# Ratio of hydrophobic-hydrophilic and positive-negative residues at lipid-water-interface influences surface expression and channel-gating of TRPV1

**DOI:** 10.1101/2020.09.04.272484

**Authors:** Somdatta Saha, Rashmita Das, Divyanshi Divyanshi, Nikhil Tiwari, Ankit Tiwari, Ritesh Dalai, Abhishek Kumar, Chandan Goswami

**Affiliations:** School of Biological Sciences, National Institute of Science Education and Research, Jatni Campus, Bhubaneswar 752050, Orissa, India; Homi Bhabha National Institute, Training School Complex, Anushakti Nagar, Mumbai 400094, India; Institute of Bioinformatics, International Technology Park, Bangalore, 560066 India; Manipal Academy of Higher Education (MAHE), Manipal 576104, Karnataka, India

**Keywords:** Lipid-water interface, Capsaicin, channelopathy, Ca^2+^-influx, membrane damage, Dye uptake

## Abstract

During evolution, TRPV1 has lost, retained or selected certain residues at Lipid-Water-Interface (LWI) and formed specific patterns there. The ratio of “hydrophobic-hydrophilic” and “positive-negative charged” residues at the inner LWI remains conserved throughout vertebrate evolution and play important role in regulating TRPV1 trafficking, localization and functions. Arg575 is an important residue as Arg575Asp mutant has reduced Capsaicin-sensitivity, surface expression, colocalization with lipid-raft markers, cell area, and increased cell lethality. This lethality is due to the disruption of the ratio between positive-negative charges there. Such lethality can be rescued by either using TRPV1-specfic inhibitor 5’-IRTX or by restoring the positive-negative charge ratio at that position, i.e. by introducing Asp576Arg mutation in Arg575Asp backbone. We propose that Arg575Asp mutant confers TRPV1 in a “constitutive-open-like” condition. These findings have broader implication in understanding the molecular basis of thermo-gating and channel-gating and the microenvironments involved in such process that goes erratic in different diseases.

## Introduction

Ion channels have unique abilities to sense different physical and chemical stimuli and conduct influx of specific ions. Due to these unique abilities, ion channels are involved in plethora of complex cellular functions including regulation of volume and ionic homeostasis, signal transduction as well as propagation of action-potential^1^. Notably, function/s of ion channels are heavily dependent on the membrane environment, i.e. change of pH, voltage difference across membrane, binding of a ligand, temperature, and also by the composition of lipid bilayers, the physico-chemical properties, surface charges etc.^2,3,4,5,6^. Never-the-less, within the lipid bilayer, ion channels undergo quick conformational changes and this conformational switching between closed and open states is referred to as channel-gating^7^. However, the complex interplay of these different stimuli, the molecular mechanism of quick conformational changes in ion channels in response to these stimuli, and how such factors contribute in the channel functions are not well understood. However, slight alterations in such complex system leads to non-functionality of ion channels leading to pathological conditions and diseases commonly known as channelopathies^8^. Escalation in the number of channelopathies has triggered the need to understand structure-function relationship of ion channels, their gating mechanisms, influence of membrane microenvironments in channel functions and finally importance of ion channels in in physiology and pathophysiology. Such understanding will enable designing of new drugs and/or effective gene therapy for treating such abnormalities^9^.

TRPV1, also known as the “Capsaicin receptor” is the finding member of TRPV subfamily and has been well characterized in the context of different physiological and sensory functions^10^. TRPV1 act as a polymodal channel which gets activated by a plethora of exogenous and endogenous stimuli like capsaicin, different endogenous compounds, low pH, temperature>43°C, etc.^11^. Interestingly, TRPV1 knock-out animals (TRPV1^-/-^) are fertile and survive without major problems suggesting that TRPV1 is not an essential ion channel *per se*^12,13^. In human population, TRPV1 have large number of genetic variations^14^. Yet, so far no disease or channelopathy has been linked with the point-mutations of TRPV1^12^. At least one copy of TRPV1 gene is present in all vertebrates and in different tissues ranging from haploid gametes to peripheral sensory neurons^15,16^. Therefore, involvement of TRPV1 in sensory and physiological functions are most likely to provide certain adaptive and physiological benefits to the individuals^17^. For this reason, molecular changes in TRPV1 sequence during vertebrate evolution can be correlated well with its structure and function/s. Such changes or no changes in the amino acids in key positions of TRPV1 can provide important clue for channel functions in different membrane micro-environments that differ in different organisms, physiological conditions and also ecological niches.

TRPV1 also represents a unique member of only few ion channels that have thermo-gating behaviour. Recently high-resolution structure of TRPV1 in open- and close-conformation has been resolved^18^. Yet, how thermo-gating works and what are the factors that regulates thermo-gating behaviour is poorly understood^19^. It is suggested by several studies that membrane components regulate TRPV1 channel function to a large extent. Indeed, importance of different lipids, PIP2, DAG as well as cholesterol in the regulation of channel function has been reported^6^. Interestingly, inverse relationship of cholesterol in TRPV1 channel function has been suggested by several groups^20^. Previous reports suggest that cholesterol forms bond with Arg residues present in the inner lipid-water-interface region in close conformation but not in open conformation^21^.

The availability of TRPV1 sequences from diverse species, the high-resolution structure of TRPV1 in open- and closed-conformation, human genome data offers unique opportunity to analyse the importance of individual amino acids in channel functions, especially the amino acids present in the lipid-water interface region, i.e. approximately 6-10 Å thin-layer of microenvironment present on both sides of the lipid bilayer. This work suggests that TRPV1 has evolved with unique combinations of amino acid that maintain a critical ratio of hydrophobic-hydrophilic nature. We propose that the conserved ratio of positive-negative charged amino acids on the inner LWI region of TRPV1 is a critical parameter that regulates the channels’ surface expression as well as function.

## Results

### Analysis of molecular selection and exclusion of amino acids in the lipid-water interface regions of TRPV1 throughout vertebrate evolution

A total of 37 mammals, 7 birds, 3 reptilians, 3 amphibians and 4 piscean TRPV1 sequences were considered. The frequency-of-occurrence of all 20 different amino acids at the lipid-water-interface (LWI) region was calculated. The same calculation was also performed for the outer as well as inner LWI region separately (**Fig 1**). Analysis reveals that the frequency-of-occurrence of positively-charged amino acids in inner LWI remain mostly constant during vertebrate evolution, however with different values. Amongst all the positively-charged amino acids, Arg (having the highest pI of 13.5 for the side group) occurs most frequently (10%) in the inner LWI region of TRPV1 and this percentage remain conserved from amphibians to mammals. The other positively-charged amino acids, Lys (pI = 10.4 for the side group) also show a conserved frequency-of-occurrence (6.67%) across evolution on the inner leaflet, especially from amphibians to mammals. However, frequency-of-occurrence of another positively-charged amino acid, namely His (pI = 6.8 for the side group) remain conserved, yet at a very low value or at zero value suggesting that His residue is never selected in the inner LWI region while other positively-charged residues were selected there. Notably, the frequency of Arg, Lys and His in the inner LWI region also suggests that positively-charged residues with high pI values (for side group) only are selected and retained in the inner LWI region (discussed later).

**Fig 1.**
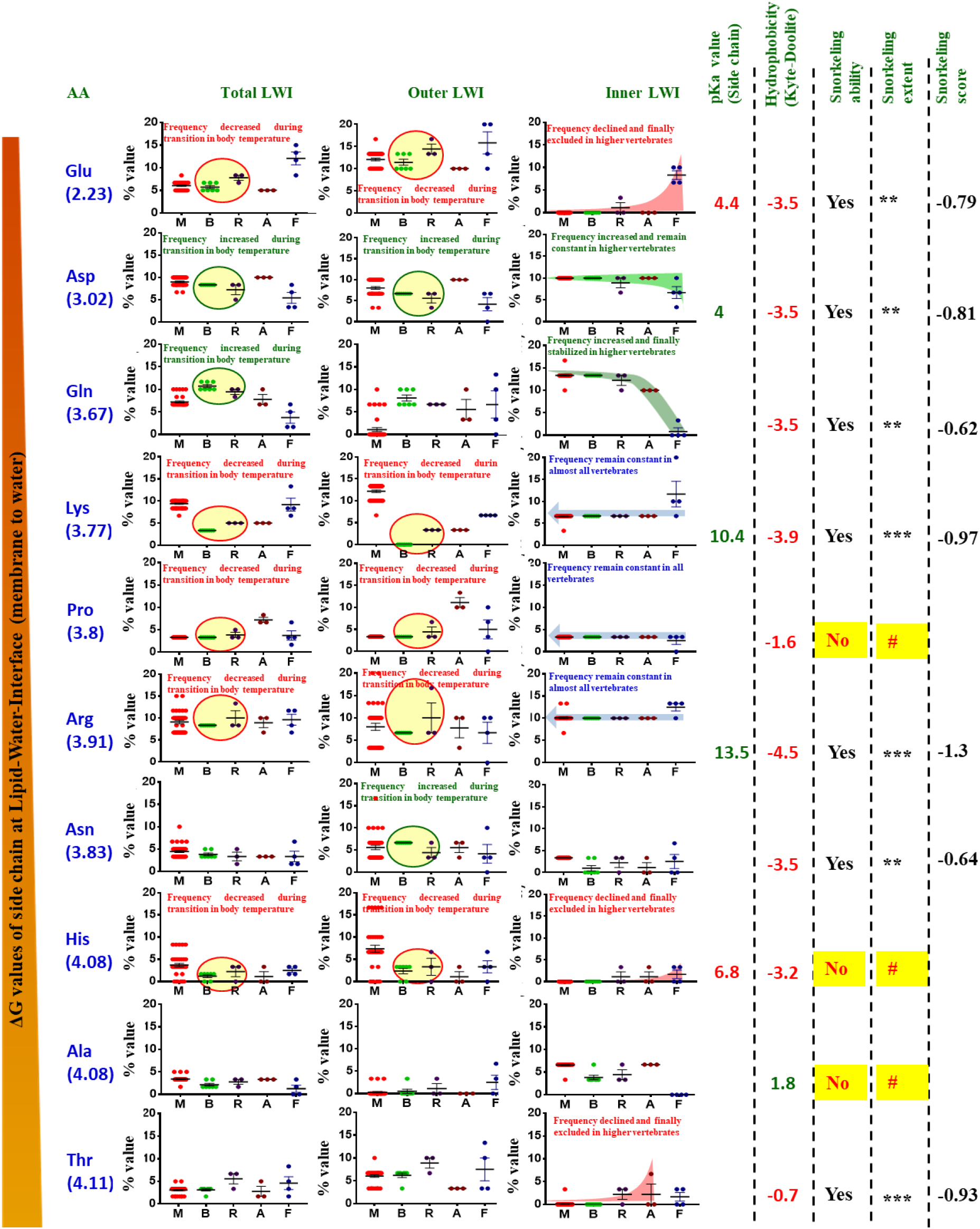
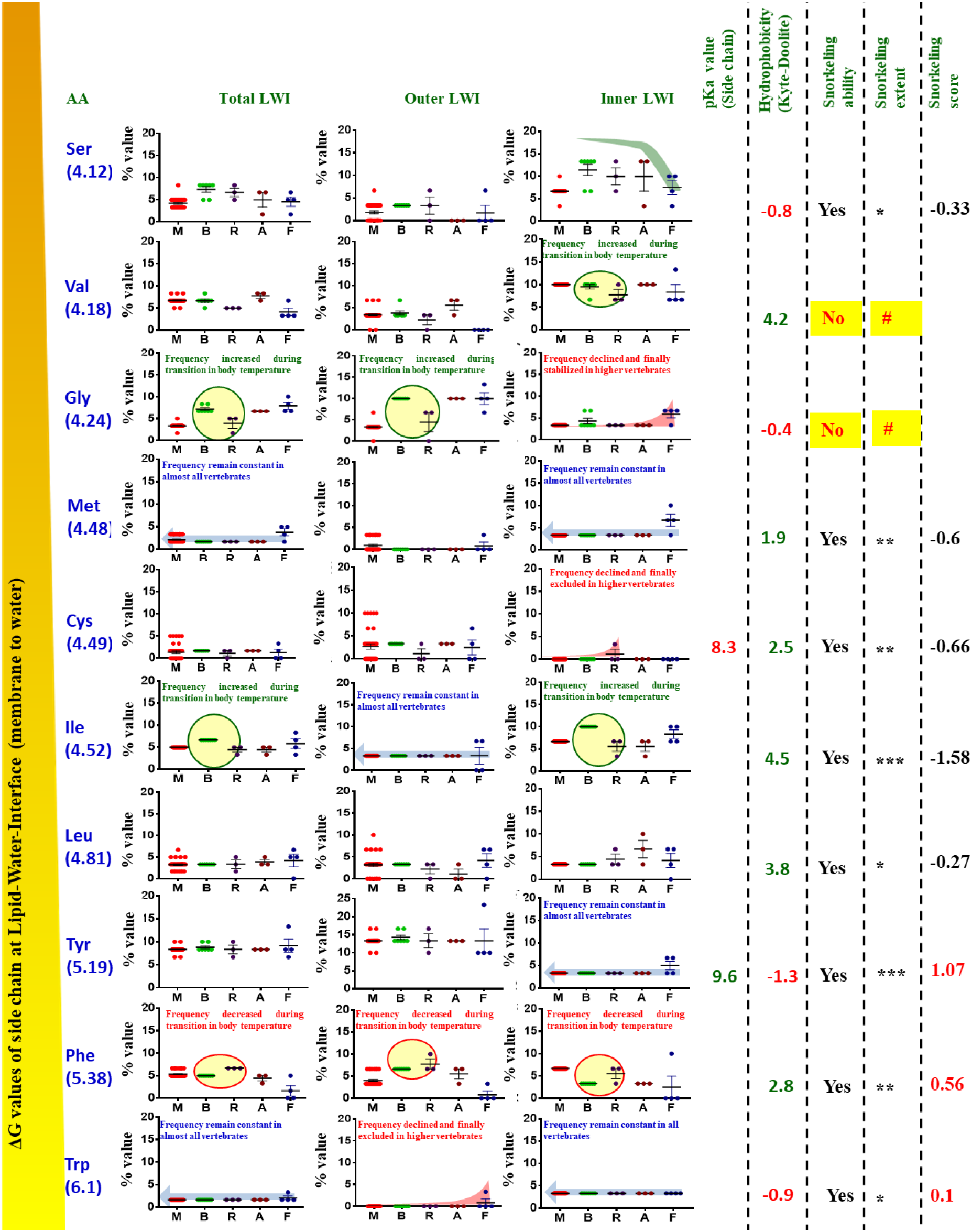
“Frequency-of-occurrence” analysis for different amino acids at the lipid-water-interface (LWI) of TRPV1 in different phylogenetic groups. The “frequency-of-occurrence” of all 20 different amino acids at inner LWI (right-most row), at outer LWI (middle row), and total (combining inner and outer LWI, left-most row) have been shown separately. Frequency-of-occurrence of different amino acids of TRPV1 LWI region across different vertebrate species ranging from fishes (F), amphibians (A), reptiles (R), birds (B) and mammals (M) are shown. The amino acids were arranged according to the low to high ΔG values of the side-chain values^22^. In cases where percentage of amino acids remain constant are indicated by blue arrow as back ground. Similarly, positively selected and negatively selected (or excluded) are marked with green or red background. In right side, the snorkelling extent of each amino acid is represented in *** (high), ** (moderate), * (low) and not at all (#). The circles with green (for positive selection) or red (for negative selection) lines indicate the changes in frequency-of-occurrence during transition from reptiles to birds as a result of change in body temperature.

For negatively-charged amino acids we observed a reverse pattern of occurrence, especially at the inner LWI region. For the negatively-charged residues at the inner LWI region, higher the pI, lower is the frequency-of-occurrence. Asp residue (pI of 4 for the side group) occurs more frequently on the inner leaflet as compared to Glu residue (pI of 4.4, for the side group). Moreover, the % frequency of Asp residue on the inner LWI has increased from fishes to amphibians (indicative of positive selection) and subsequently remained conserved throughout the evolution (indicative of stabilization). On the other hand, frequency of Glu has decreased from nearly 10% in fishes to almost 0% in amphibians (indicative of negative selection), and remain at 0% value to mammals suggesting gradual exclusion of this amino acid in this LWI region. However, in the outer LWI region, frequency-of-occurrence of Glu and Asp residues are random and remain at a relatively high frequency.

Aromatic amino acids, namely Tryptophan, Tyrosine and Phenylalanine have relatively low frequency-of-occurrence across evolution in the inner LWI. Both Tryptophan and Tyrosine remain at less than 5% frequency, yet remain conserved at the inner LWI region throughout the vertebrate evolution (indicative of critical function played by these amino acids). Frequency-of-occurrence of Phenylalanine remain random in the inner as well as outer LWI region. Out of these three aromatic amino acids, only tyrosine (pI of 9.6 for the side group) retains a higher frequency-of-occurrence (13%) at the outer LWI.

Helix breaking amino acid Proline has almost equal distribution throughout evolution both in outer (3.33%) and inner LWI (3.33%) region, especially in the higher vertebrates. Another helix-breaking amino acid, namely Gly has been retained at less than 5% frequency at the inner LWI region, but mostly remain conserved throughout the vertebrate evolution. Individual hydrophobic amino acids, namely Isoleucine, Leucine, Valine and Alanine retain different percentages of frequency-of-occurrence and conservation. Notably, Isoleucine remains conserved in outer LWI. Among other amino acids Met remains conserved in the inner LWI region. At inner LWI region, the frequency-of-occurrence for Cys is low in all vertebrates and eventually excluded in the higher vertebrates. Taken together, the analysis suggest selection, retention or exclusion of specific amino acids at the inner LWI region and therefore provides important clue of TRPV1 function in different species (discussed later)

### Ratio of positive-negative amino acids in the inner LWI region remain constant in TRPV1 through-out the vertebrate evolution

All positively-charged amino acids (Arg, Lys, His) have a higher frequency-of-occurrence at the outer LWI region in mammals but its pattern of occurrence is not conserved throughout evolution. Though the frequency-of-occurrence of positively-charged amino acids is less on the inner LWI but its frequency-of-occurrence has remained conserved throughout vertebrate evolution from amphibians to mammals (**Fig 2a**). Similarly, negatively-charged amino acids have a higher frequency-of-occurrences on the outer LWI, but is more conserved on the inner LWI. While frequency-of-occurrence for individual amino acids are variable, the ratio of the total number of positively-charged amino acids and total number of negatively-charged amino acids at the inner LWI show a ratio of 1.67:1 and this ratio remain conserved in the entire vertebrate evolution. This ratio is not conserved in case of outer LWI. These findings strongly suggest that during vertebrate evolution, this conserved ratio at the inner LWI has played a strong selection pressure in case TRPV1 function (discussed later).

**Fig 2.**
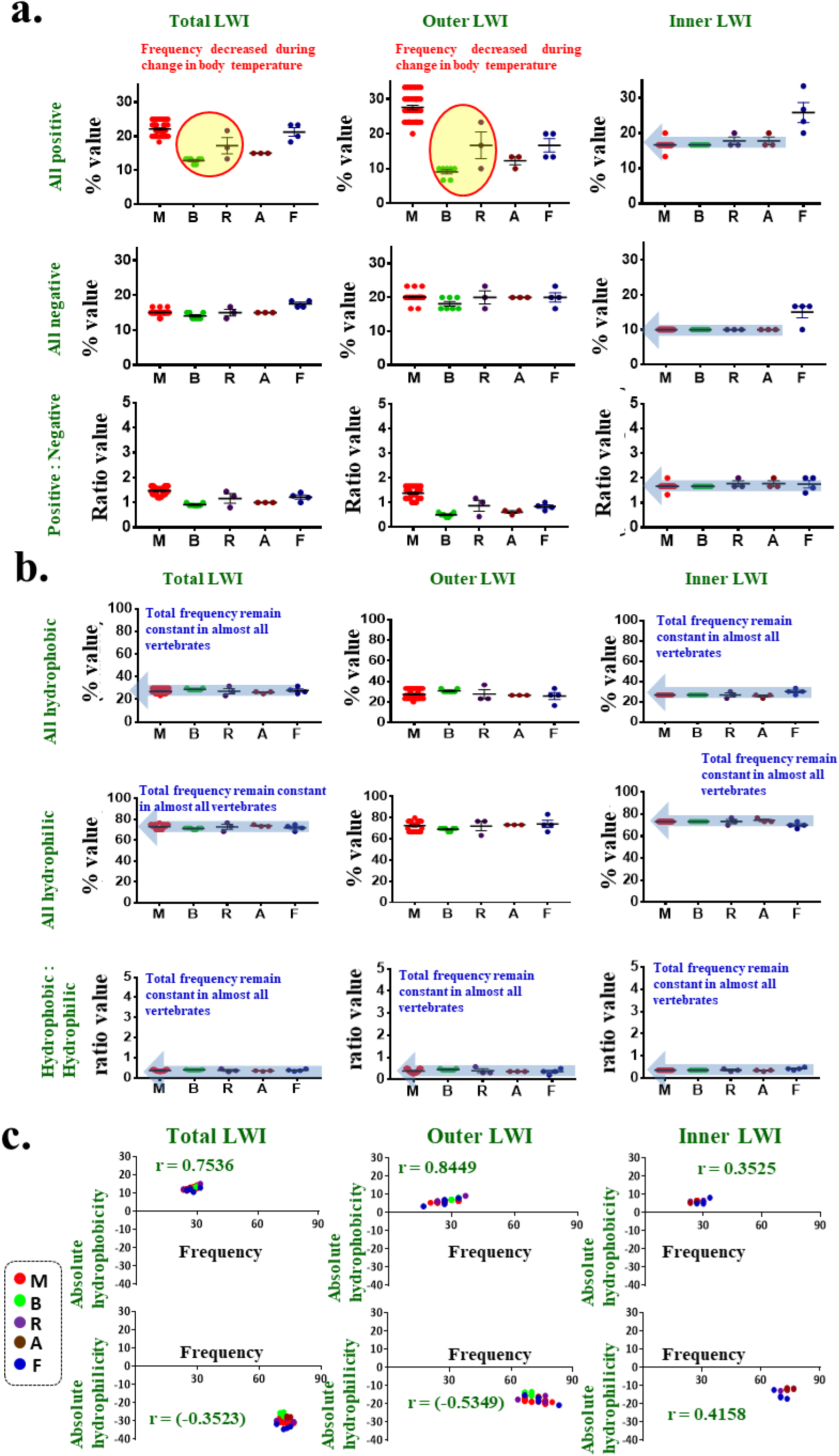
Ratio of positive/negative amino acids and ratio of hydrophobic/hydrophilic amino acids remain constant at the inner lipid-water-interface of TRPV1. Frequency-of-occurrence of (a) positively- and negatively-charged residues and (b) hydrophobic and hydrophilic amino acids at the lipid water interface of TRPV1 during vertebrate evolution are shown. **a.** Positively-charged amino acids (Arginine, Lysine, Histidine) and negatively-charged amino acids (Aspartic Acid and Glutamic Acid) at the lipid water interface region of TRPV1 across different vertebrate species ranging from fishes, amphibians, reptiles, birds and mammals has been calculated. Negatively-charged amino acids on the inner leaflet appear to be more conserved across evolution. The ratio of positively-charged to negatively-charged amino acids on the inner leaflet at a value of 1.67:1 remain conserved throughout vertebrate evolution. **b.** The frequency-of-occurrence of hydrophobic amino acids (Cys, Phe, Ile, Leu, Met, Trp, Tyr), hydrophilic amino acids (Ala, Asp, Glu, Gly, His, Lys, Asn, Pro, Gln, Arg, Ser, Thr, Val) and their ratio are shown The ratio of hydrobhobic/ hydrophilic/ amino acids at the inner leaflet remain conserved at a value of 0.36:1 throughout the vertebrate evolution. **c.** The frequency (plotted in the X-axis) of hydrophobic or hydrophilic amino acids correlates with absolute hydrophobicity or hydrophobicity (plotted in Y-axis) at the LWI regions. The values for Pearson correlation coefficient (r) are provided along with the graphs.

### TRPV1 retains unique combination of hydrophobic and hydrophilic amino acids in its inner lipid-water interface

Hydrophilic amino acids (Ala, Asp, Glu, Gly, His, Lys, Asn, Pro, Gln, Arg, Ser, Thr, Val) have a higher incidence of occurrence (~73.14% in total) on the inner LWI region as compared to the hydrophobic residues (Cys, Phe, Ile, Leu, Met, Trp, Tyr, Total ~26.85%). This difference in distribution of hydrophobic and hydrophilic amino acids at the inner LWI region has been more or less conserved across evolution from fishes to mammals. However, when the ratio of occurrence of the all hydrophobic and all hydrophilic amino acids is considered, the ratio remain conserved in the inner, outer and total LWI (**Fig 2b**). These findings strongly suggest that during vertebrate evolution, the conserved ratio of hydrophilic versus hydrophobic amino acids at the LWI regions of TRPV1 has also played a strong selection pressure in case TRPV1 function (discussed later).

As different amino acids have different hydrophobicity index, we explored if “frequency” can reflect reliably for the “absolute total hydrophobicity or hydrophilicity”. For that purpose, we calculated the absolute hydrophobicity or hydrophilicity in each species and plotted the values. The analysis suggests a fairly good conservation in the total hydrophobicity or hydrophilicity values (considering the side chains and peptide bond contribution group of the amino acids as mentioned in in total, outer and also in inner LWI regions ^22^ (**Fig 2c**). The absolute hydrophobicity or hydrophilicity values correlate well with the frequency-of-occurrence of these amino acids also (**Fig 2c)**. These also suggests that the frequency-of-occurrence in the LWI region can be reliably used as a bio-physical parameters relevant for the structure-function relationship.

Combination of amino acids offering glycosylation or phosphorylation do not show any trend, neither in inner LWI nor in outer LWI regions (**Fig S1**). Therefore, the data clearly suggest that ratio of hydrophilic-hydrophobic amino acids as well as ratio of positive-negative charge plays important selection pressure, at least in the inner LWI region.

### TRPV1 is localized in the cholesterol-enriched lipid rafts and Arg575Asp mutation induce cell lethality

TRPV1 in closed conformation interacts with cholesterol mediated by the Arginine residues present at 557 and 575 positions^21^. Substitution of Arginine by a negatively (Aspartic Acid) or neutrally charged (Alanine) amino acid resulted in altered localization of TRPV1 in lipid rafts. In this work, we explored if positive charge due to presence of Arginine in these positions are critical or if substitution of these Arg with any other positively-charged residue like Lys or His is capable of retaining same properties. For this reason, we generated 4 more substitution mutants: Arg557His, Arg557Lys, Arg575His and Arg575Lys. All these mutations (we term all these as LWI mutants) were made on rTRPV1-WT-GFP, i.e. Arg557Lys, Arg557His, Arg575Lys, Arg575His, Arg557Ala, Arg557Asp, Arg575Ala and Arg575Asp were subsequently transfected in DRG neuron derived F-11 cell line to analyse the localization of these mutants with respect to TRPV1-WT-GFP (**Fig 3**). To check the lipid raft localization of rTRPV1-WT-GFP and these LWI mutants, the transfected cells were stained with CTXB-594 under normal as well as in cholesterol reduced conditions (MβCD, 5mM treated conditions) (**Fig 4**).

**Fig 3.**
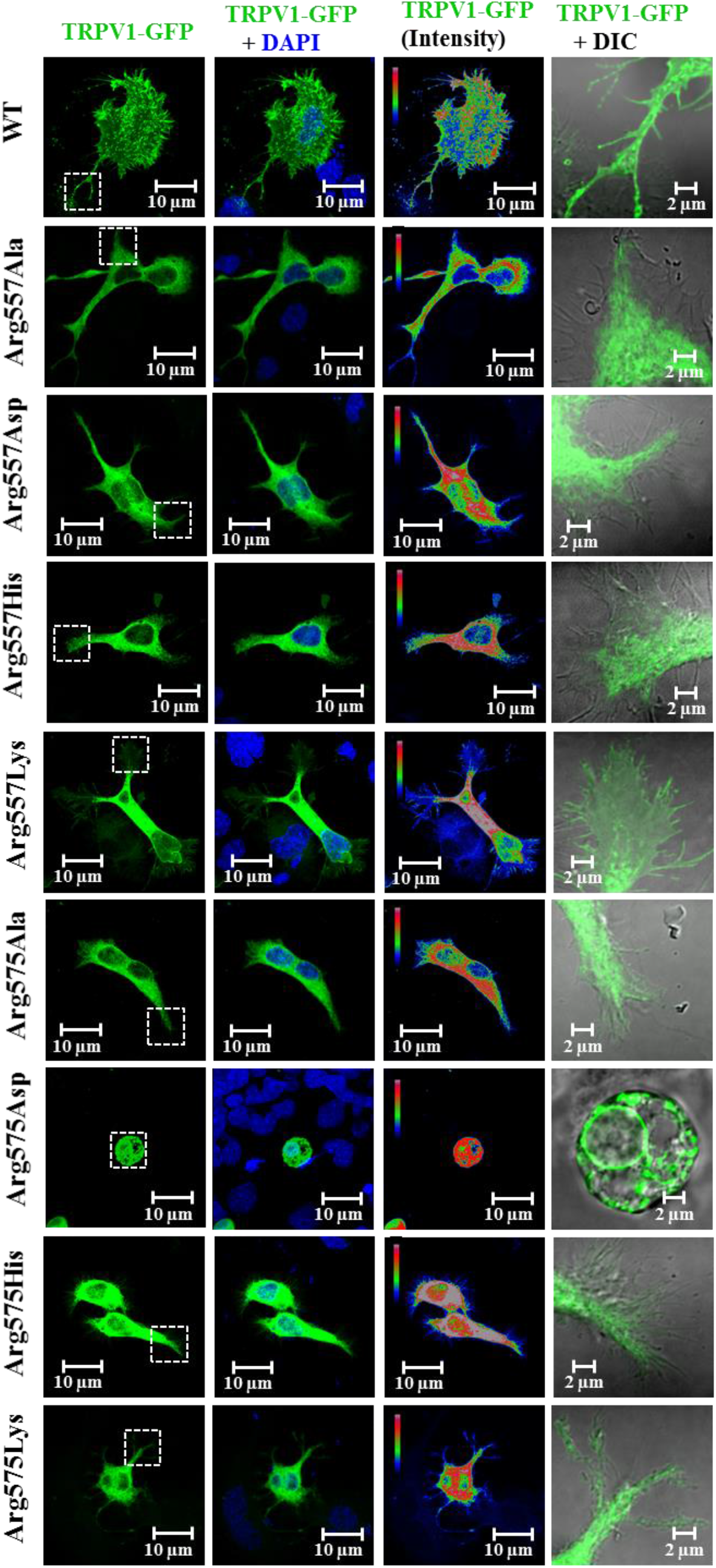
Arg residues at the Lipid-Water-Interface (LWI) of TRPV1 are required for its proper surface expression and membrane localization. The localization pattern of rTRPV1-WT-GFP or different LWI mutants in GFP has been shown. These GFP-tagged proteins (green) were transiently expressed in F-11 cells for 36 hours, fixed and imaged using confocal imaging. WT-rTRPV1 shows distinct membrane localization, whereas the LWI mutants fail to localize at the membrane. Often the LWI mutants are retained in the ER and/or cause fragmentation of ER. The intensity of GFP-tagged proteins are shown in the rainbow scale. Nucleus is stained with DAPI (blue). Enlarged view of surface areas (indicated by dotted square) for GFP fluorescence at the membrane merged with DIC image are shown in the right side.

**Fig 4.**
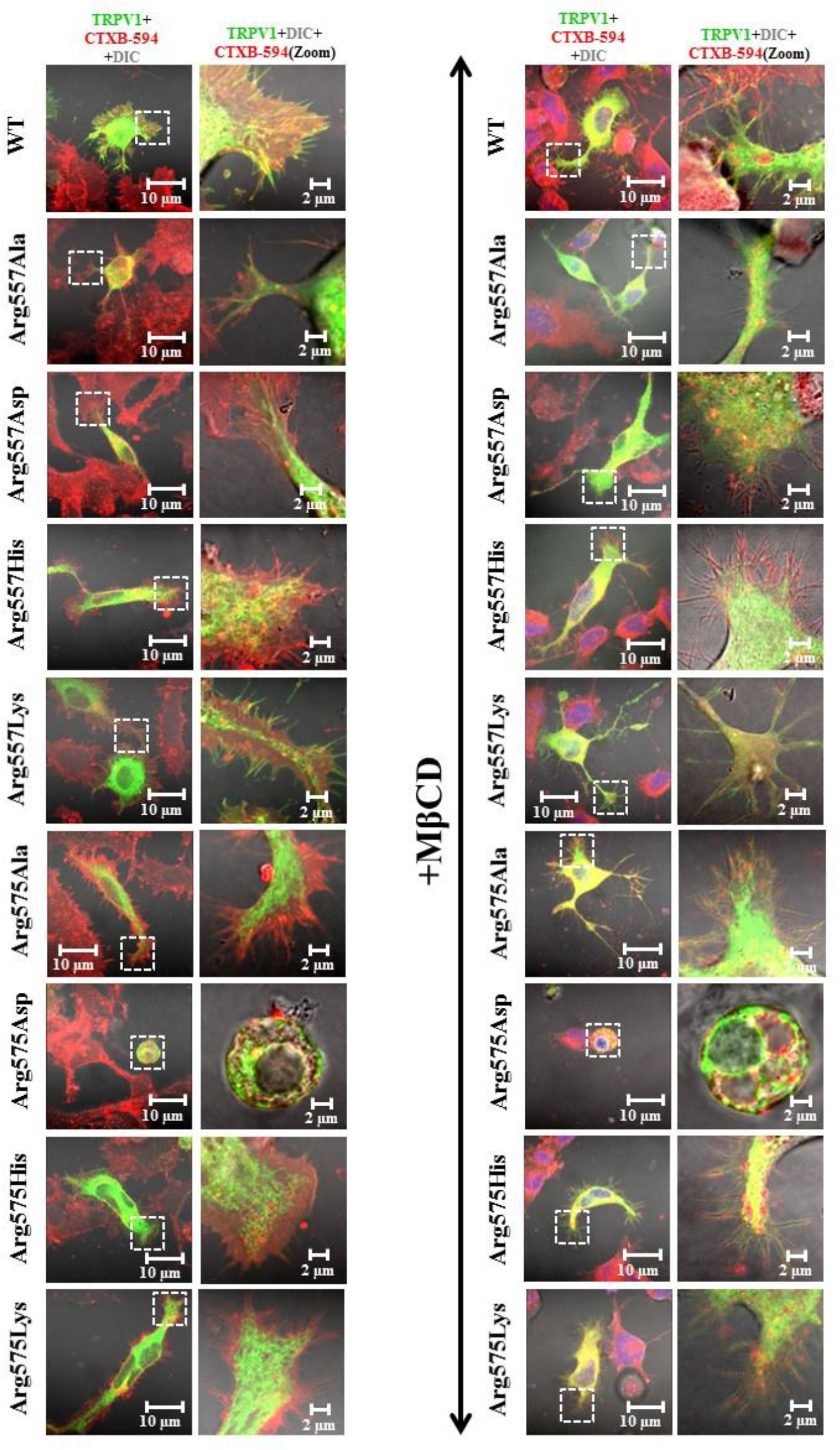
TRPV1-WT but not the LWI mutants co-localize with endogenous lipid raft markers. TRPV1-WT-GFP (green) and different LWI mutants in GFP (green) were expressed in F11 cells. Cells were fixed 36 hours post transfection and stained for lipid raft with Cholera Toxin B-594 (red). All images were acquired by confocal microscope. Same experiment was performed with cells that were treated with 5mM MβCD for cholesterol depletion 30 min before cell fixation. TRPV1-WT-GFP shows distinct co-localization with Cholera Toxin B in the membranous region while LWI mutants are distinctly excluded from Cholera Toxin B-enriched membranous regions.

Unlike TRPV1-WT, none of these LWI mutants are capable of localizing to the plasma membrane or to the ends of the neurites and/or filopodia (**Fig 3**). Compared to TRPV1-WT which co-localize with the lipid raft markers, the mutants fail to show co-localization under control or even cholesterol-depleted conditions (in MβCD-treated conditions). TRPV1-WT show no co-localization with CTXB in MβCD-treated conditions. In an over-expression system, when these GFP-tagged constructs were co-transfected along with two other lipid raft markers Flotillin-1-RFP and Caveolin-1-RFP (**Fig 5 and Fig S2**), we obtained similar results. Caveolin1 and Flotillin 1 co-localized with TRPV1-WT-GFP but not with the LWI mutants which show little or no co-localization. Thus, Arginine at positions 557 and 575 is important for proper localization of TRPV1-WT to the lipid rafts. All the TRPV1-LWI mutants exhibit membrane trafficking problems, yet none of these are lethal for the cells except for Arg575Asp mutant. Most of the F-11 cells expressing TRPV1-Arg575Asp tend to assume a rounded morphology soon after transfection (in ~ 24 hours), exhibit no neurites or filopodial structures and were often observed to detach from the surface on which they were grown. The Arg575Asp mutant mostly localized to the ER membrane. We conclude that TRPV1-Arg575Asp mutant imparts lethality to cells expressing them.

**Fig 5.**
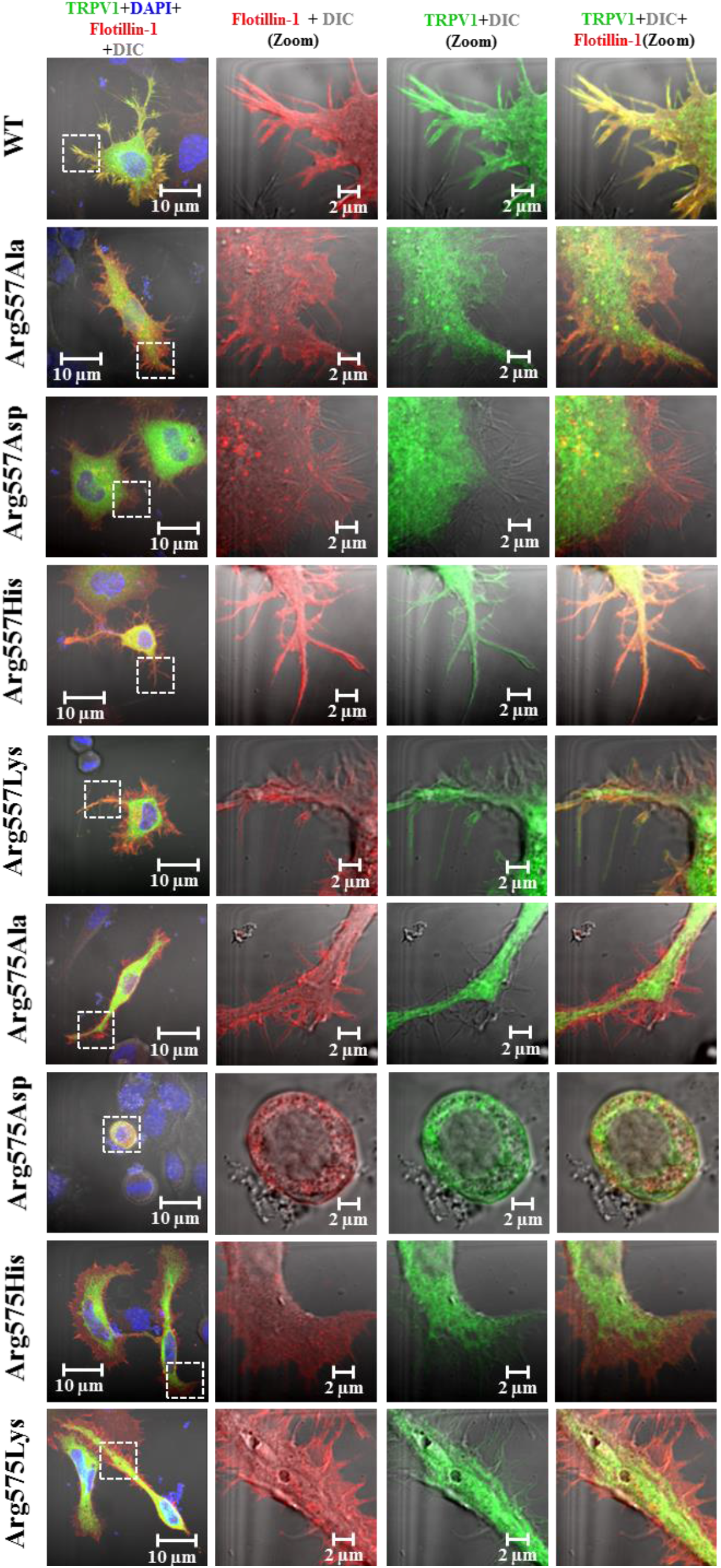
TRPV1-WT but not the Lipid Water Interface (LWI) mutants co-localize with overexpressed lipid raft markers. GFP-tagged (green) TRPV1 wild type (WT) and different LWI mutants were co-expressed with lipid raft marker Flotillin1-RFP (red) in F11 cells. Cells were fixed 36 hours post transfection and images were acquired by confocal microscope. TRPV1-WT shows distinct co-localization with Flotillin1-RFP in the membranous region. Notably, the LWI mutants are distinctly excluded from Flotillin1-RFP enriched membranous regions even after over expressing both.

### Arg575Asp lethality can be rescued by maintaining overall positive-negative charge ratio

As this Arg575Asp mutation disrupts the positive-negative charge ratio at the inner LWI (especially in the TM5 region which regulates pore opening), this perturbation could be one of the plausible cause leading to constitutive channel opening and thus lethality observed in the cell. In order to explore this possibility, we mutated the adjacent residue having a conserved Aspartic Acid at 576 position to Arginine in the TRPV1-Arg575Asp mutant. Notably, the ratio of positive- and negatively-charged residue in this double mutant is same as in wild-type TRPV1. Interestingly, F11 cells expressing the double mutant, i.e. TRPV1-Arg575Asp/Asp-576Arg-GFP retain their normal morphology. Therefore, the lethality observed in case of TRPV1-Arg575Asp can be rescued by introducing positive charge at 576^th^ position (**Fig 6a**). When the length, breadth, area and perimeter of F-11 cells transfected with TRPV1-WT-GFP, TRPV1-Arg575Asp-GFP and TRPV1-Arg575Asp/Asp576Arg-GFP were calculated, we found a significant increase in the length, perimeter and area of cells transfected with TRPV1-Arg575Asp/Asp576Arg-GFP as compared to cells expressing TRPV1-Arg575Asp-GFP (**Fig 6b**). However, even though TRPV1-Arg575Asp/Asp576Arg-GFP was capable of retaining the viability of transfected cells, its migration to the plasma membrane and its capability to form neurites and filopodia remained impaired in comparison to TRPV1-WT-GFP. TRPV1-Arg575Asp/Asp576Arg-GFP was neither able to co-localize with overexpressed lipid raft markers like Caveolin1-RFP and Flotillin1-RFP, nor with lipid rafts endogenously stained with CTXB-594 in normal as well as cholesterol reduced conditions (**Fig S3**). This suggests that TRPV1 is optimized for a positive charge at 575^th^ position and negative charge at 576^th^ position. This particular phenotype attributed by TRPV1-Arg575Asp was not specific to F-11 cells. In SaOS and HaCaT cells too, the cells attained rounded morphology when TRPV1-Arg575Asp was expressed. This can be effectively rescued by TRPV1-Arg575Asp/Asp576Arg double mutant. This data indicates that the phenotype observed is independent of cell types (**Fig S4**).

**Fig 6.**
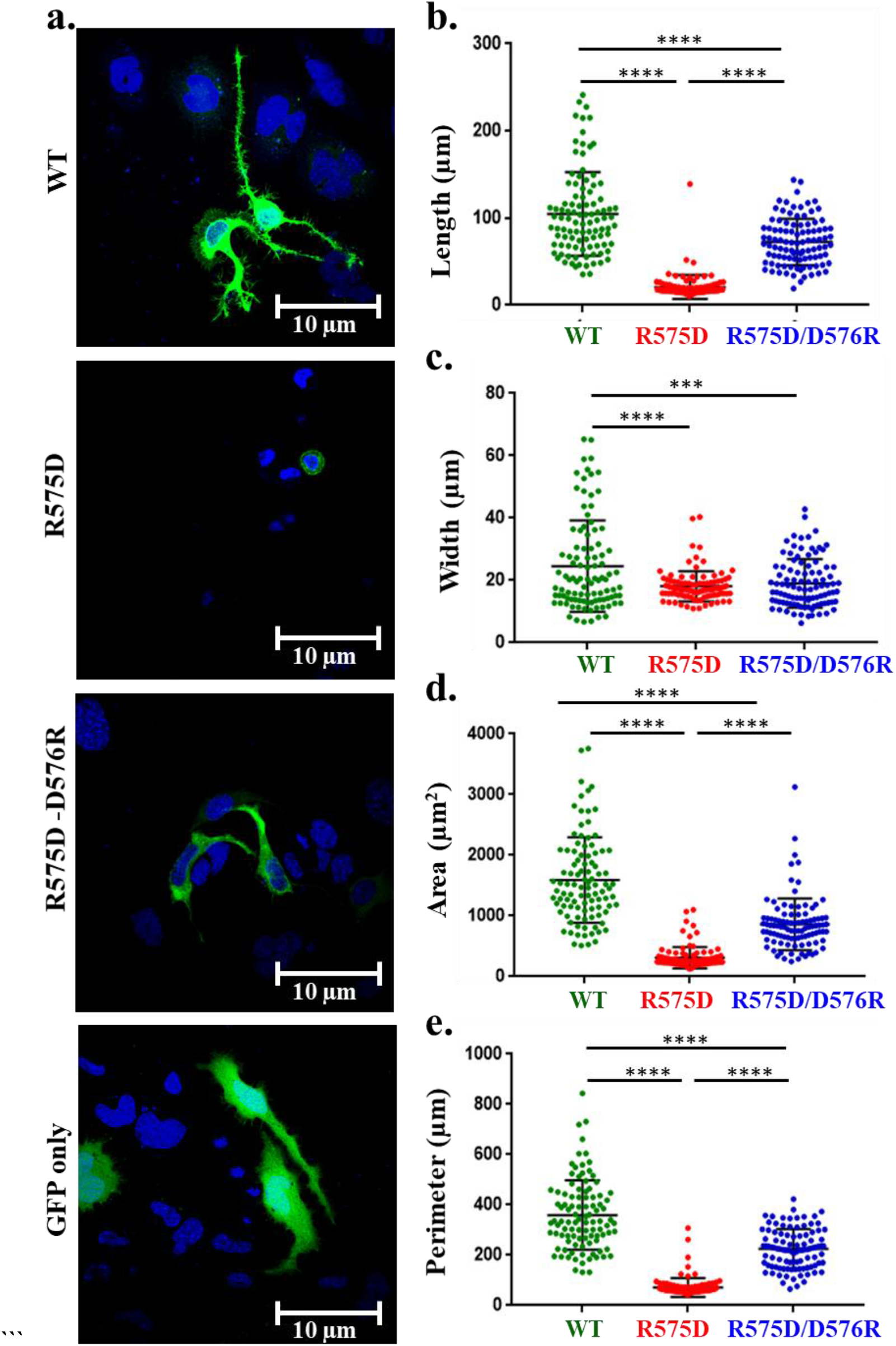
Cell lethality due to Arg575Asp mutation can be rescued by introducing another mutation. **a.** Confocal images of TRPV1-WT, TRPV1-Arg575Asp and TRPV1-Arg575Asp/Asp576Arg in GFP vector expressed in F11 cells. Cells expressing Arg575Asp mutation tend to assume a circular morphology and decreased surface adhesion behaviour. **b.** Quantification of different morphology parameters such as length, width, area and perimeter suggest that Arg575Asp mutant induce reduction in cell size and introduction of Asp576Arg on Arg575Asp rescues all these parameters. In all cases, minimum 100 cells were quantified. The **** (P <0.0001) and *** (P<0.0003) indicate values that are significantly different.

### Cell lethality due to TRPV1-Arg575Asp can be partly rescued by long-term channel inhibition

We hypothesized that one of the reasons for Arg575Asp lethality might be constitutive channel opening. In order to test this hypothesis F-11 cells were transiently transfected with TRPV1-WT-GFP and TRPV1-Arg575Asp-GFP, treated with 1μM 5’-IRTX for 36 hours and then fixed with 4% PFA. Images of treated as well as untreated transfected cells were acquired and the length, breadth, area, perimeter of cells for each condition was measured. Constitutive application of TRPV1 specific channel blocker 5’-IRTX prevented lethality of cells expressing Arg575Asp suggesting that the cell lethality is mainly due to the excess channel function *per se* (**Fig 7**).

**Fig 7.**
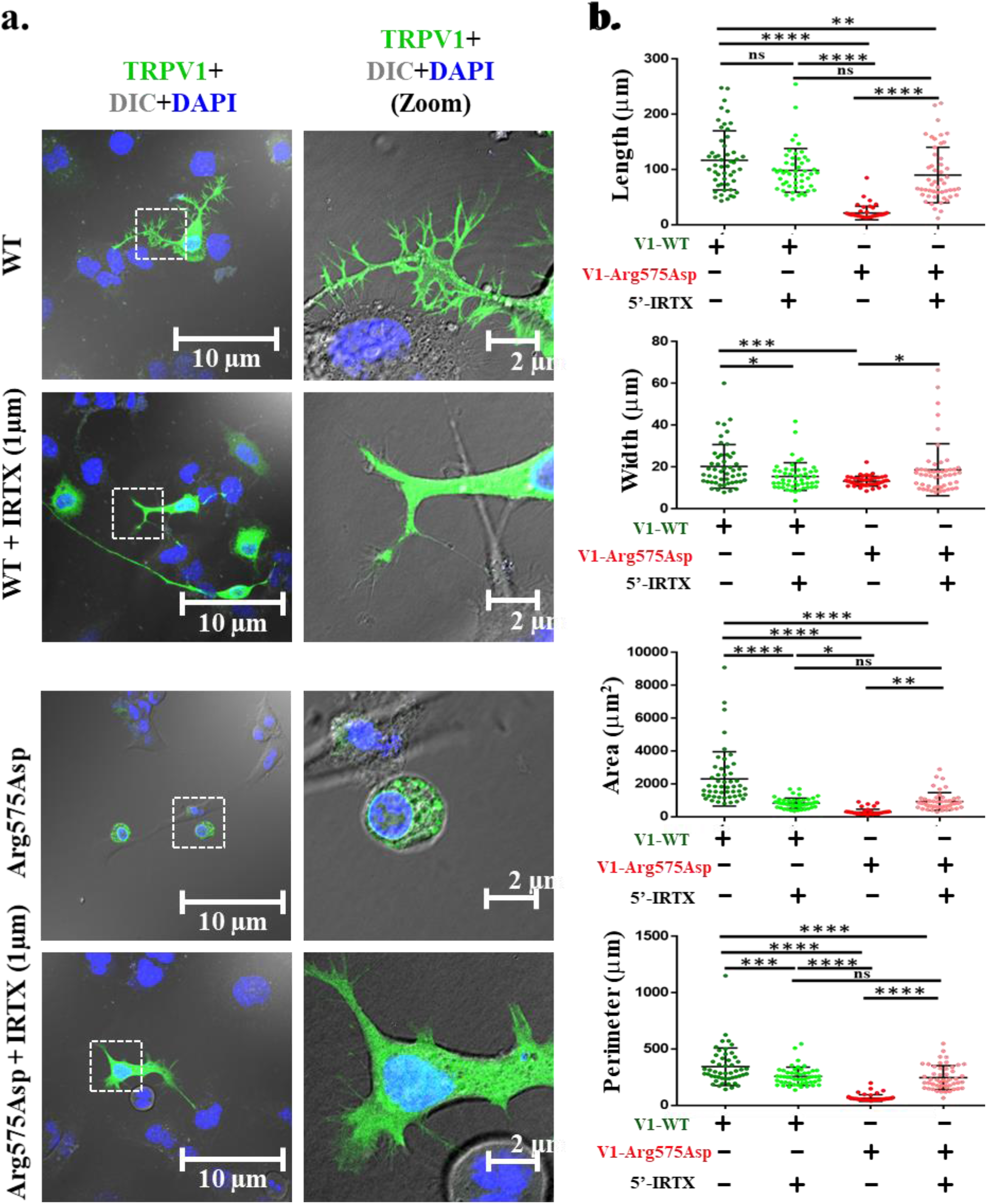
Cell lethality due to Arg575Asp mutation can be rescued by TRPV1 channel blocker. **a.** Confocal images of TRPV1-WT-GFP and TRPV1-Arg575Asp-GFP expressed in F11 cells and grown in absence and presence of 5’-IRTX, a specific inhibitor of TRPV1. F11 cells expressing TRPV1-Arg575Asp-GFP become much elongated and produce neurites and filopodia in presence but not in the absence of 5’-IRTX. **b.** Quantification of different morphology parameters such as length, width, area and perimeter of cells mentioned above. In all cases, minimum 50 cells for each condition were quantified. The ** (P = 0.0098), **** (P<0.0001) and *** (P=0.0006) indicate values that are significantly different while ns indicate values that are non-significantly different.

### Arg575 and Arg579 are located close proximity of Asp576

Next we explored the distribution of Arg575 residue in TRPV1 in close and open conformation. For that purpose, we used the closed and open structure of TRPV1 and also made homology model of Arg575Asp mutation. The results suggest that Arg575, Arg579 and Asp576 residues are located in a cluster and in close proximity to each other in slightly upper side of the inner lipid water interface (**Fig 8**). These residues show minor change in positioning within this cluster in close as well as in open conformation. In case of Arg575Asp mutation, these three residues are still located in the same cluster, yet results in change in ratio of the charge distribution within this cluster. This suggests that such close-proximity and charge-sharing might be relevant for channel opening. As Arg575 residue is located close to the lower gate, it also suggests that Arg575Asp mutation probably fix the channel with a bigger pore or a “open-like channel”.

**Fig 8.**
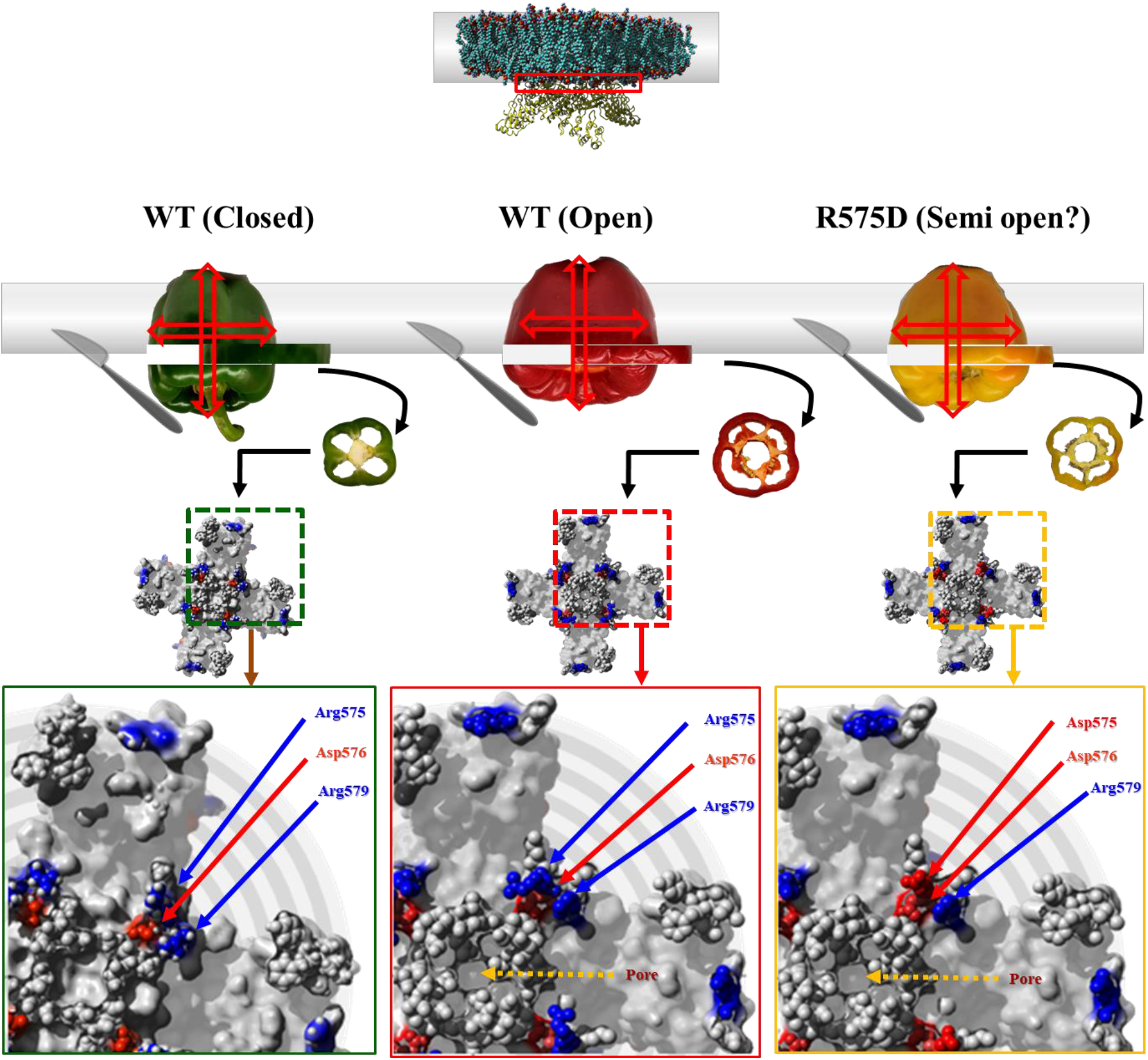
Positive and negative charged amino acids share close proximity at the inner Lipid-water-interface region. Shown are the structure of a thin-layer of TRPV1 representing wild-type TRPV1 in close and open conditions as well as Arg575Asp mutant in open conformation. Positive charges of Arg575 and Arg579 remain located in close proximity to Asp576. The nature of the ion channel and the position of the layer is schematically shown by using capsicum of different colour and its cut versions.

### Cells expressing TRPV1-WT and TRPV1-Arg575Asp mutation uptakes more Rhod-3 AM dye

TRPV1 is known to induce enhanced uptake of FM464 dye in F11 cells^23^. TRPV1 is known to have larger pore size that can allow much larger molecules that are otherwise impermeable through the cell membrane. For example, TRPV1 pore allow QX-314 (a charged molecule with a molecular mass of 263 Da), 6-Methoxy-N-ethylquinolinium iodide (commonly known as MEQ, a cell membrane impermeable Choline indicator dye of molecular mass 315.15), FM1-43 (a cell impermeable dye of molecular mass 452 Da), and YO-PRO (Cell membrane impermeable dye with a molecular weight:629.3216) to pass through TRPV1 pore in cells that express TRPV1^24,25,26,27^. TRPV1 pore is also permeable to N-Methyl-D-glucamine (molecular weight of 195.21)^27^.

In this work, we used Rhod-3 AM dye (molecular weight ~1600 Da) loading efficiency as a tool to probe the membrane permeability of the non-transfected F11 cells and cells expressing TRPV1-WT and mutants. We noted that Rhod-3 fluorescence is much less in non-transfected cells as well as in GFP expressing cells. However, Rhod-3 fluorescence is much more in TRPV1-WT expressing cells compared to non-transfected cells, as visualized in the same view field. Even significant fluorescence signal is achieved in TRPV1-WT cells when less Rhod-3 dye is incubated for less time and imaged in reduced laser powers. This data strongly suggests that Rhod-3 permeability as well as entrapment within F11 cell is not same in all cells and is enhanced significantly by the presence of TRPV1-WT. The higher level of Rhod-3 fluorescence signal within the cell is achieved in Arg575Asp mutant, but at higher concentration of dye and at a longer time of incubation, suggesting that the mutant may cause some difficulty in the dye uptake and/or dye entrapment within the cell as compared to the wild type TRPV1. However, Rhod-3 fluorescence signal is very low in TRPV1-Arg575Asp/Asp576Arg mutant. Taken together this data suggests that TRPV1-Arg575Asp mutation probably induces large pores in cells expressing them whereas Arg575Asp/Asp576Arg double-mutant does not support uptake of Rhod-3 AM dye (**Fig S5**).

### TRPV1-Arg575Asp mutant forms a Capsaicin-insensitive channel that is rescued by Arg575Asp/Asp576Arg

F-11 cells were transiently co-transfected with full-length Rat TRPV1-WT (present in pmCherryC1 vector) and a Ca^2+^-sensor pGP-CMV-GCaMP6f. The same was repeated for TRPV1-Arg575Asp (present in-pmCherryC1 vector), TRPV1-Arg575Asp-Asp576Arg (present in-pmCherryC1) and only mCherry (empty vector of pmCherryC1). Approximately 24 hours post transfection, live cell imaging of only doubly transfected cells were executed using Olympus FV3000 confocal microscope for 200 frames (time gap between two frames is 1.085 seconds). Capsaicin (10 μM) was added at the 30^th^ frame for each condition. Unlike, TRPV1-WT which showed a rapid increase in Ca^2+^-influx upon Capsaicin addition, TRPV1-Arg575Asp showed no visible change in fluorescence intensity upon Capsaicin application. In order to establish the fact that the imaged cells are live and functional at the time of experiment, Ionomycin (2μM) was added almost at the end (120^th^ frame), which resulted in the formation of a Ca^2+^-spike. TRPV1-Arg575Asp/Asp576Arg exhibited similar Capsaicin-sensitivity like that of TRPV1-WT. However, capsaicin, failed to show any Ca^2+^-influx in cells transiently transfected with only pmCherryC1 and TRPV1-Arg575Asp-pmCherryC1. Quantification of pGP-CMV-GCaMP6f fluorescence intensities from doubly transfected cells show that both TRPV1-WT and TRPV1-Arg575Asp-Asp576Arg are “Capsaicin-responsive” and thus a peak is immediately formed after the 30^th^ frame and another one at the 120^th^ frame upon addition of Ionomycin which opened all Ca^2+^-channels. TRPV1-Arg575Asp and pmCherryC1 remained unresponsive to Capsaicin and show a peak only upon addition of Ionomycin after the 120^th^ frame (**Fig 9**).

**Fig 9.**
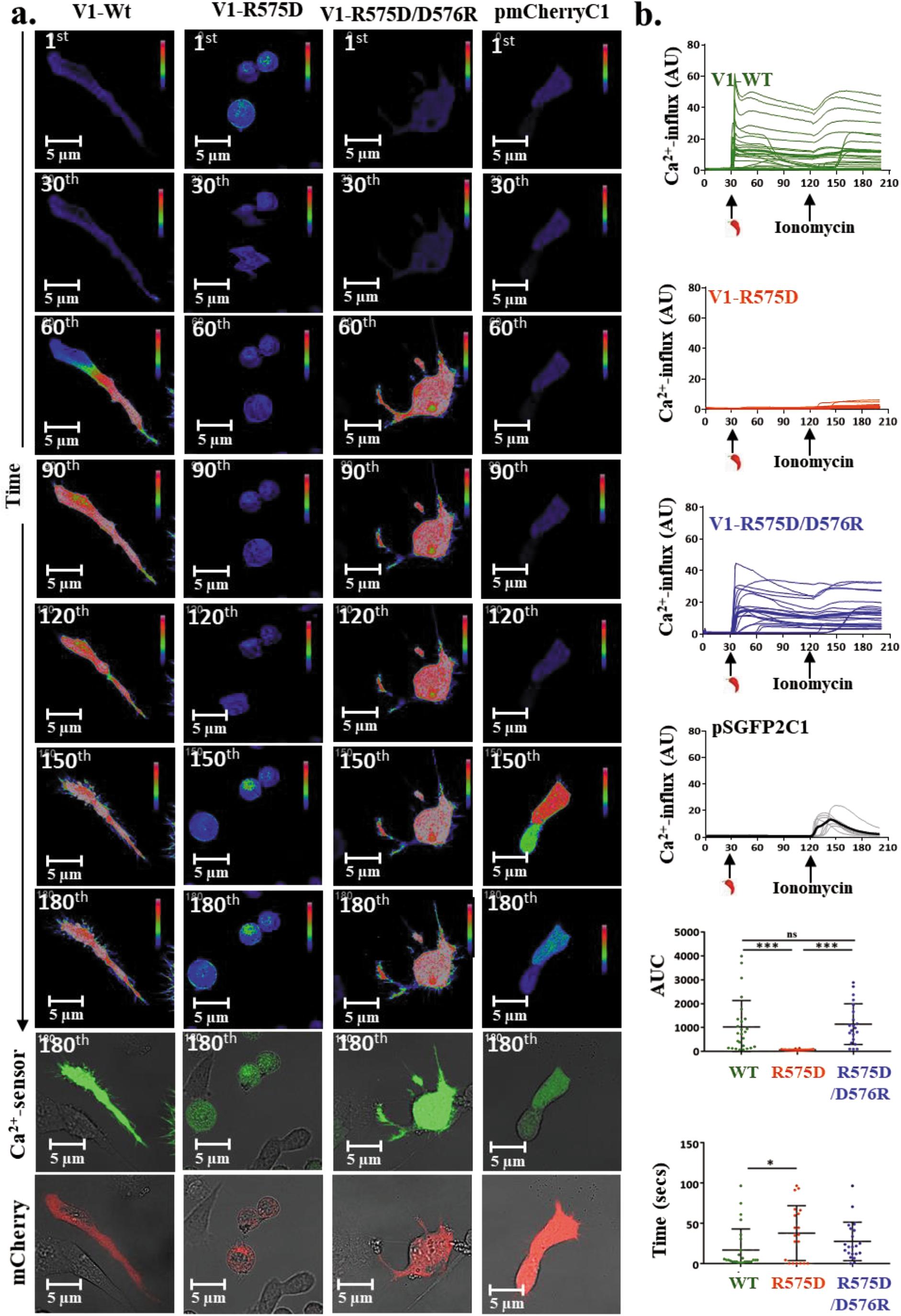
Ca^2+^-imaging of TRPV1-WT, Arg575Asp, Arg575Asp-Asp576Arg and pmCherryC1. F-11 cells doubly transfected with TRPV1-WT-pmCherryC1, TRPV1-Arg575Asp-pmCherryC1, TRPV1-Arg575Asp/Asp576Arg-pmCherryC1 and pmCherryC1 and the Ca^2+^-sensor pGP-CMV-GCaMP6f shows levels of Ca^2+^-influx upon addition of TRPV1 specific agonist Capsaicin (10 μM) at 30^th^ frame and the ionophore Ionomycin (2 μM) at 120^th^ frame. **a.** TRPV1-WT and TRPV1-Arg575Asp-Asp576Arg show a rapid influx of Ca^2+^ upon Capsaicin activation. The change in fluorescence intensity of pGP-CMV-GCaMP6f before and after agonist stimulation has been depicted in RGB mode. TRPV1-Arg575Asp remained insensitive to (P.T.O) Capsaicin addition. **b.** Quantification of pGP-CMV-GCaMP6f fluorescence intensity from cells transfected with TRPV1-WT (n = 26 cells), TRPV1-Arg575Asp (n=20 cells), TRPV1-Arg575Asp/Asp576Arg (n = 21 cells) and pmCherryC1 (n = 8 cells) shows a graphical representation of the change in intensities upon addition of Capsaicin at the 30^th^ frame and Ionomycin at the 120^th^ frame. The quantification for all the constructs were done considering the initial value recorded at the 1^st^ frame as 1. The intensities for the remaining frames were subsequently calculated relative to the 1^st^ frame. The thick line in each intensity plot demarcates the average intensity of each construct. For each individual cell, Area Under Curve (AUC) was calculated for 1-120 frame and Time (required to reach the point of highest intensity) was calculated for 30-120 frame.

## Discussion

In this work we analysed the conservation of different amino acids in the lipid-water interface of TRPV1 and evaluated the importance of critical residues. Such understandings provide critical information about functionality in different ion channels which are of importance.

### Mutation-induced channelopathies and channel gating

So far several diseases have been identified that are linked with the dysfunction of ion channels, collectively termed as “channelopathies”. The most common cause of such impairment is mutation in genes encoding ion channels. Such mutations affect different properties such as channel folding, proper localization, relevant interactions with lipids and/or proteins, post-translational modifications, and often channel-gating behaviour^28^. As TRPV1 is a non-selective cation channel and polymodal in nature (i.e. it gets activated by different chemicals and temperature), changes in TRPV1 sequence is indicative of suitability of amino acids in micro-environments at the lipid-bilayer suitable for proper channel gating. As TRPV1 is involved in thermo-sensory functions and other physiological functions, improper channel gating, i.e. “constitutive-open” or “constitutive-close” nature of mutations is expected to provide either disadvantages or at least provide no adaptive advantages. Therefore, uncontrolled spontaneous opening of TRPV1 is expected to cause lethality at the level of cells, tissues and also in individual. These results confirm that TRPV1 retains a specific ratio of hydrophobic-hydrophilic residues (0.36:1) as well as positive-negative charged residues (1.67:1) in its inner LWI region throughout vertebrate evolution. In spite of the variation in “frequency-of-occurrence” in individual amino acids, the conservation in these ratios are highly significant and shades important information about the structure-function relationship of TRPV1. Notably, these values remain constant in all vertebrates. In fact, there is no change in these ratio values in reptiles as well as in birds (indicating transition from cold-blooded animals to warm blooded animals), and the same values exists in early vertebrates too suggesting that these values actually indicate functions that are independent of body temperature. We propose that these ratio values indicate suitability of amino-acids in specialized micro-environments required for “channel-gating” and not for “thermo-gating” *per se*.

### Importance of membrane compositions in the channel functions

Importance of different membrane compositions in channel localization and function is well established. In this context, it is important to mention that TRPV1 activity is modulated by various exogenous and endogenous lipid molecules^29^. For example, mutations that abolish TRPV1-PIP2 interaction results in TRPV1 having lower chemical and thermal activation thresholds^30^. In other word, PIP2 helps TRPV1 to stabilize in a closed-state. Depletion of membrane cholesterol by MβCD-treatment results in decreased expression and activity of TRPV1 in Rat DRG neuronal membrane^31^. Cholesterol depletion also affects the pore-size and dilation of TRPV1^27^ This accords well with studies demonstrating that TRPV1 is localized in cholesterol-enriched lipid rafts and that its channel properties are altered by membrane cholesterol^32^. In fact TRPV1 has at-least one cholesterol-binding site in the S5-helix and interaction of TRPV1-cholesterol results in inhibition of channel opening^33^. This aspect matches well with other TRPV-induced channelopathies as well. For example, mutation (R616Q) causing loss-of-interaction with cholesterol can results in formation of constitutive open TRPV4 channel which cause physiological disorders^34^. Previously it has been shown that Arginine residues present at inner LWI, especially at 557 and 575 positions of TRPV1 form bonds with cholesterol in the closed but not in open conformation of TRPV1^21^. Accordingly, substitution of Arg with any other neutral (Ala) or negatively charged (Asp) amino acid resulted in lack of TRPV1 localization at membranes and with lipid-raft markers^21^. Inhibition of spontaneous opening of TRPV1 by membrane components seem to be logical as TRPV1 has a relatively large pore which is permeable to several different ions (including Ca^2+^) and even large molecules^35,24,25,26,27^. This fact is supported by the fact that under hypocalcemic conditions, persistent agonist application induces a time-dependent dilation of TRPV1 pore and thereby allowing the penetration of large cations^27^. However, under the same conditions reduction of cellular cholesterol by ~54% inhibits such pore dilation of TRPV1 and its subsequent permeability to large cations like NMDG^27^. Taken together, it is apparent that cholesterol and other membrane components play important regulatory function relevant for TRPV1 localization and is also involved in channel gating property, though the underlying mechanisms are not clear. In this context, it is important to note that selection of amino acids in the LWI regions and also in the TM regions may suggest a coevolution process influenced by the presence of certain membrane components, such as cholesterol^36^.

### LWI region of TRPV1 and importance of positively charged amino acids

Lipid-water-interface, provides a special microenvironment on both-sides of the membrane where bulk water level is low^37^. Notably, the protonation and deportation of amino acid side chains are expected to be different in LWI regions than what is considered in standard aqueous solution and thus pI values of titratable side groups of amino acids are different in LWI regions^22^. In many cases, the trans-membrane helices of the integral membrane proteins consist of apolar amino acids flanked by charged amino acids at the lipid-water-interface region which allow stable insertion of these TMs in the lipid bilayer^38^. These interfacial regions are often demarcated by charged residues (like Lysine, Arginine), or other snorkeling amino acids (such as Tyrosine and Tryptophan). Among positively-charged residues, Arginine and Lysine have better “snorkeling” ability depending on the availability of the free water in such LWI regions^39^. The long-charged side chains of Arginine and Lysine can stretch themselves to place the charged moiety in the polar lipid-water interfacial regions while keeping the hydrophobic part of their side chains in the hydrophobic core of the lipid bilayer. This kind of arrangement has been observed in several other ion channels and other proteins too. Membrane-inserted Arginine residues has been found in several ion channels (such as in case of Mg^2+^-channels)^40^, anti-microbial peptides^41^, pore-forming peptides^42^ and in case of cell-penetrating peptides too^43^. Insertion of the most hydrophilic amino acid Arginine (pKA of the side group 13.5) into the highly hydrophobic core requires sufficient amount of energy. However, it has been suggested that the guanidinium group present in the side chain of Arginine is capable of forming as many as six hydrogen bonds and hence its occurrence in a membrane protein is mainly localized to the interfacial region where it can interact with both water molecules and polar lipid moieties^44^. Considering that surface of lipid bilayer is enriched with negative charge (due to head groups of phospholipids), positioning of highly positively charged residues in the LWI region fits well there, especially for a purpose of minimizing the lateral movement of TM regions. The distributions of positively charged residues at LWI regions is often coupled with the specific arrangements of lipid/cholesterol-interacting motifs suitable for proper stabilization of the TM regions in lipid bilayers and also for the thermodynamic events experienced during conformational changes required for channel opening^45^.

### Importance of decoding the amino acid distribution in LWI

The frequency-of-occurrence of Arg, Lys, Pro, Trp and Tyr, remain constant in inner LWI throughout the vertebrate evolution. Notably, Cys amino acid is completely excluded in inner LWI throughout the vertebrate evolution, suggesting that Cys residue is misfit there. Residues such as His, Glu, and Thr gradually declined and ultimately excluded in mammals, suggesting that these amino acids are not suitable in the inner LWI regions, especially in warm blooded animals with respect to the channel functions. While positive charged amino acids such as Arg and Lys are selected at high-frequency, exclusion of His in the inner LWI is highly suggestive. High-frequency of Arg and Lys accords well with their ability to snorkel as well as their ability to form bonds with membrane components^46,47^. For example, Arg and Lys are able to form bonds with the −OH group of cholesterol at the LWI region, while His residue is unable to form such bonds. Also, considering the protonation-deprotonation possibilities in physiological pH at the inner LWI, His appears to be a misfit amino acid there^48^. Reduction in frequencies of Phe, Asn, Ala, Leu, and increment in frequencies of Ile, Val, Gln in the LWI correlates with the transition from cold-blooded animals to warm blooded animals (reptiles to birds). Therefore, these changes are more-likely to accommodate the bio-physical changes that took place in lipid membrane at low body temperature to warm temperature. The frequency-of-occurrence of helix breaking amino acids, namely Proline and Glycine remain conserved throughout vertebrate evolution, especially in the inner LWI region. The frequency-of-occurrence of hydrophobic and hydrophilic amino acids correlates well with actual hydrophobic-hydrophilic values (of the side groups) at the LWI regions. Aromatic amino acids such as Trp and Tyr remain conserved in the inner LWI region throughout vertebrate evolution. Other aromatic amino acid, namely Phe is positively selected in mammals in inner LWI. In this context, it is important to mention that the aromatic residues determine the preferred rotational and dynamics of membrane-spanning segments relevant for the function of membrane proteins^49^.

### What we learn from the frequency calculation: importance of certain amino acids in channel functions

Our analysis sheds important information about the structure-function relationship of TRPV1. Notably, in inner LWI region, amino acids with titratable side group has followed three distinct trajectories. Either such amino acids were retained at a conserved frequency, such as Asp or Lys residues in the inner LWI region, or such amino acids were gradually excluded. For example, Glu residues have been gradually excluded from the inner LWI regions. On the other hand, some amino acids with titratable side group, such as Arg have been positively selected. From our data it become also evident that the amino acids that have retained a high frequency-of-occurrence (~10%, and at constant values) during vertebrate evolution in the inner LWI are Arginine and Aspartic Acid. In fact, Arg and Asp share 1:1 ratio at the inner LWI region, especially in higher vertebrates. Data suggest that Arg at 575^th^ position may play important role in the channel gating, possibly by altering the conformation of TM5 and thus the lower-gate of TRPV1^50^. It seems that Arg575 is involved in the interaction of other negatively charged residues in order to channel opening and thus involved in channel gating. For example, Arg575 can form salt-bridges with Glu692 at physiological temperature^51^. Similarly, Asp576 forms salt bridge with Thr685 at physiological temperature and also at higher temperature with as a result of heat deactivation^51^. Based on the molecular simulation data, charge neutralization of Arg575, Arg579 and Lys694 by PIP2 has also been suggested^52^. Based on the sequence conservation and experimental evidence, we propose that it is possible that at physiological pH, the net charge at the inner LWI of TM5 is neutral [due to presence of a positive (Arg575) and negative charge (Asp576)] and ability of the Arg residue to interact with cholesterol may allow TM5 to be fixed in a conformation that is equivalent to “closed state”, yet flexible enough for opening in presence of proper stimuli (**Fig 10**). Considering that the inner surface of the lipid bilayer is intrinsically negatively charged (due to the presence of phospho-groups), when the inner LWI region of the TM5 is fixed with more negative charge, the TM5 may shift further down from the bilayer resulting in “locking of mutant channel” in either “open” or at-least “open-like” or in a “dilated” state. This is more relevant, as TRPV1 become sensitized (equivalent to more spontaneous opening) upon phosphorylation in its Ser residues, which introduces more negative charge in its loop regions^53^.

### Does Arg575Asp mutation makes it a constitutive open ion channel?

The Arg residue at 575 position, Asp residue in 576 position, and Arg residue at 579 position are actually present in a cluster and these three residues are highly conserved throughout the vertebrate evolution (**Fig S6**). Positioning of a positively charged residue followed by negatively charged residues in this stretch can be of high importance (**Fig S6**). Such arrangements have also been observed on other channels, such as in Shaker Kv channels which contain Arg followed by Glu amino acid in the inner LWI region of its TM5^54^.

Notably Arg at 575 position in TRPV1 seem to be very critical for proper localization of the channel at lipid raft and also maintaining interaction with cholesterol, at least in the closed state. As these two residues, namely Arg575 and Asp576 have not changed during the transition from reptiles (cold-blooded) to birds (warm-blooded), it can be proposed that both Arg575 and Asp576 are not involved in “thermo-gating” property *per se*, but is actually involved in “channel gating” in standard vertebrate membrane containing cholesterol. However, Arg575 is a crucial amino acid for overall channel gating and Arg575Asp mutation brings double negative charge to the end of TM5 located at LWI regions. Such strong negative charge is expected to displace TM5 further down from LWI region due to strong repulsion by negatively charged surface and lack of bond formation with membrane cholesterol may happen. Such aspects may cause “permanent locking” of Arg575Asp mutant in a constitutive open-like conformation (**Fig 10**).

**Figure 10.**
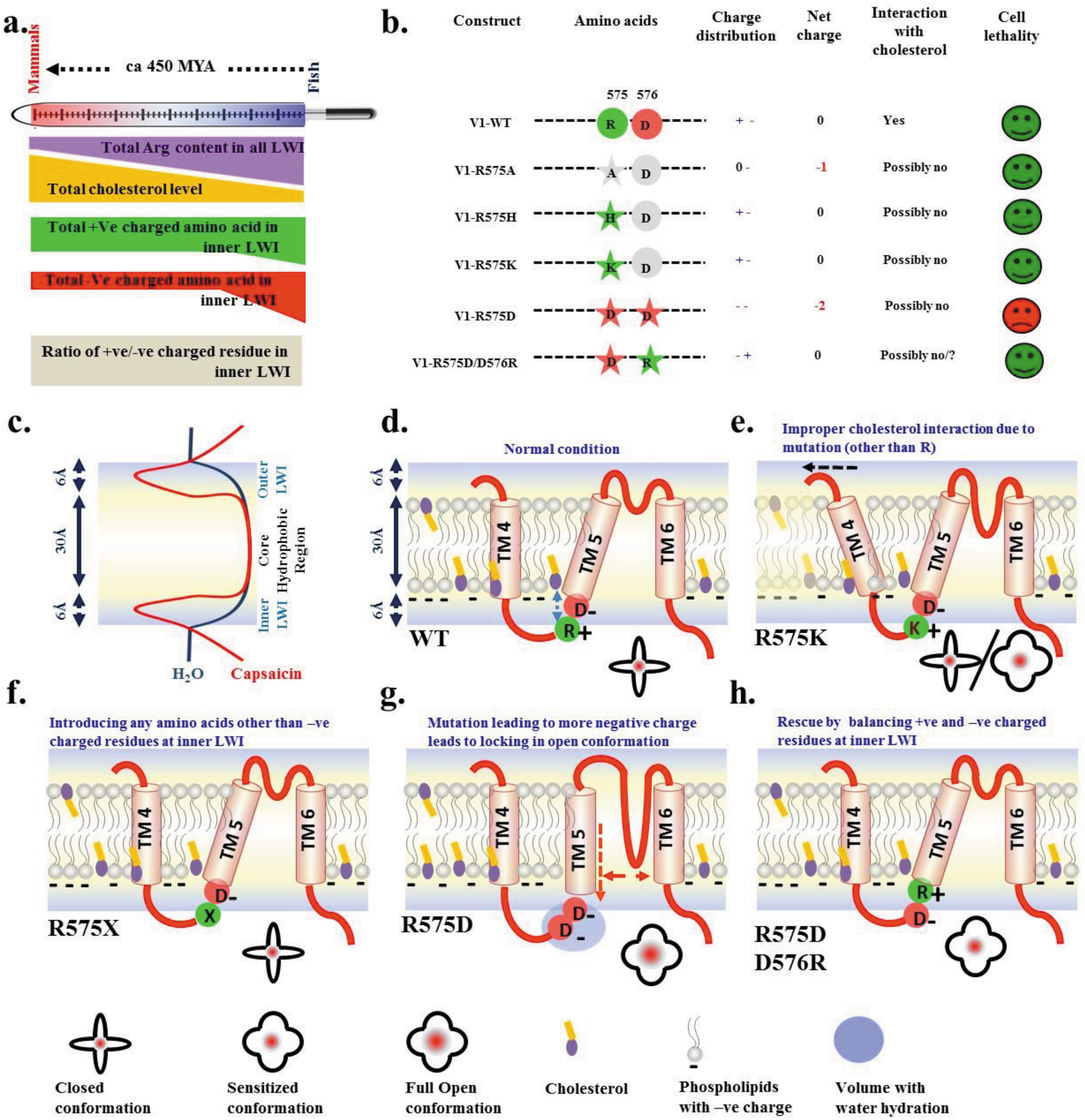
A plausible model depicting the importance of positive and negative charged amino acids in inner LWI region of TM5 in channel functions. **a.** During vertebrate evolution, average percentage of body cholesterol increases gradually while body temperature changed suddenly (warm-blooded animals evolved from cold-blooded animals). In-spite of changes in the individual values of different amino acids, the ratio of total “positively-charged” to “total negatively-charged” residue remain constant in the inner LWI. **b.** Distribution of individual charged residues as well as net charge at the inner LWI region of TM5 for wild-type and different mutations are shown. **c.** The transmembrane helices and unstructured loops are mostly occupied in the core hydrophobic lipid bilayer (~ thickness of 30Å) and both side lipid-water interface regions (~ thickness of 6A) respectively. **d-h.** Shown are the schematic representation of TM5-P-loop-TM6 of TRPV1 in different scenario. While R residues interact with −OH group of the cholesterol effectively, positive and negative charge at the inner LWI region of TM5 neutralize each other in order to position TM5 properly in an ideal confirmation suitable for channel gating. Alteration in negative-positive charge distribution there alters the positioning and conformation of TM5 as well as its possibility to interact with cholesterol. Introducing more negative charge in the inner LWI region of TM5 leads to “locking” of the TM5 in a more open-like conformation and pore dilation resulting cell lethality. Introducing a positive charge next to 575^th^ position rescue this lethality.

Indeed, there are several reasons that prompt us to propose that Arg575Asp mutation leads to a “constitutive-open like” TRPV1 channel. Firstly, expression of Arg575Asp mutant leads to cell lethality immediately (~24 hrs) after its expression in F-11 cell. The same lethality is also observed in other cell types such as SaOS and HaCaT suggesting that Arg575Asp mutation is universally lethal in different types of cells (**Fig S4**). Second, most of the cells expressing Arg575Asp mutant are insensitive to Capsaicin suggesting that a majority of the cells are already in an activated stage where further activation by Capsaicin may not be possible. Third, based on the Rhod-3 AM labelling it seems that the basal level of cytosolic Ca^2+^ is relatively high in cells expressing Arg575Asp mutation compared to other non-transfected cells. Fourth, the cells that express Arg575Asp mutant is more permeable to large compounds such as Rhod-3-AM dye when compared to other non-transfected cells. Fifth, the lethality due to Arg575Asp can be rescued by a TRPV1-specific blocker 5’-IRTX. Finally, Ca^2+^-imaging experiments suggest that even prolonged application of Capsaicin does not facilitate further entry of Ca^2+^ ions into the cells expressing Arg575Asp mutant. All these results suggest that Arg575Asp mutation induces TRPV1 to form a constitutively open channel. That possibility has been supported by the fact that introducing Asp576Arg mutation in the background of Arg575Asp rescue cell lethality, though the double mutant show defects in the lipid raft localization. Ca^2+^ imaging results suggest that Arg575Asp/Asp576Arg is capable of rescuing the channel gating function to some extent, but not the lipid raft localization. In that case non-annular interaction of TRPV1-cholesterol might be relevant.

In this context it is worth mentioning that TRPV1 is highly permeable to Ca^2+^ and it shows Ca^2+^-dependent desensitization, which effectively allows TRPV1 channel to become insensitive to stimuli when applied repetitively^55^. Notably, intracellular Ca^2+^ can interact with phosphatidylserine in physiological concentration and cause physico-chemical changes, altered hydration in the lipid-water-interface. This in turn alters lipid packing and may slow down interfacial dynamics^56^. Such changes might be useful for the Ca^2+^-dependent desensitization event of wild-type TRPV1. However, Arg575Asp mutant may have differences in that aspect.

### Conclusion

In this work we have “decoded” a molecular pattern in the lipid-water interface of the membrane protein. We also decoded the significance of certain amino acids, their patterns and their combinations in the lipid-water interface regions of TRPV1. We term this molecular pattern as “Lipid-Water-Interface Pattern Theory” (LWI-PT). We experimentally demonstrate that such pattern plays important role in channel functions and changing such code can induce cell lethality. Notably, restoring such codes can also rescue channel functions. This understanding will allow to analyse the other theromosensory and non-thermosensory channels and their functions in the light of LWI-PT. Such understanding may also help us to dissect the impact of different point mutations that induce pathophysiological disorders.

## Materials and Methods

### Generation of frequency plots

Sequences for TRPV1 were retrieved from NCBI for all Vertebrates according to previous report^21^. LWI residues were determined and percentage content of all amino acids were separately calculated using MEGA 5.1^21^. In addition to this, calculations were made for the following groups: Positively charged, negatively charged, hydrophobic and hydrophilic amino acids. In each LWI region a total 5 amino acids as a stretch was considered. Frequency calculations were done for LWI on the inside (cytoplasmic side, for 6 positions; total 30 amino acids), outside (extracellular side, for 6 positions; total 30 amino acids) as well as for the overall (i.e. for a total of 60 amino acids) residues for each species. These values were then plotted using GraphPad Prism7 (http://www.graphpad.com/).

### Calculation of absolute hydrophobicity and hydrophilicity at lipid-water-interface

Whole residue interfacial hydrophobicity scale was used for obtaining transfer free energies of each amino acid where contributions of the peptide bonds as well as sidechains were taken into consideration^22^. Whole residue scales for POPC bilayer interfaces and for n-octanol using two families of peptides: host-guest pentapeptides of the form AcWL-X-LL, for determining sidechain hydrophobicities, and the homologous series AcWLm (m = 1 to 6), for determining peptide bond hydrophobicities was used. The values used for calculating the hydrophobicity and hydrophilicity were derived from free energies of transfer of AcWL-X-LL peptides from bilayer interface to water as described before^22^. The decision level for selection of hydrophobic amino acid was taken as ΔG ? 0 and Hydrophilic as ΔG < 0.

### Site-directed mutagenesis and construct preparation

The Arg557Ala, Arg557Asp, Arg557His, Arg557Lys, Arg575Ala, Arg575Asp, Arg575His Arg575Lys mutants of TRPV1 were prepared by using site directed mutagenesis kit (Agilent Technologies) with specific primer sets as described before^21^. In all cases, full-length rTRPV1 (Rat TRPV1) cloned in pCDNA3.1 was used as a template as described earlier^21^. TRPV1-Arg575Asp/Asp576Arg mutant was generated using TRPV1-Arg575Asp as the template. In this study, Arg557Ala, Arg557Asp, Arg557His, Arg557Lys, Arg575Ala, Arg575Asp, Arg575His, Arg575Lys and Arg575Asp/Asp576Arg will be henceforth collectively referred to as LWI mutants.

After mutagenesis, full-length rTRPV1-WT and all LWI mutants were cloned into pSGFP2-C1 or GFP vector (Addgene) using 5′-CCAGGAATTCTATGGAACAACGGGCTAGC-3′ and 5′-CCAGGTCGACTTATTTCTCCCCTGGGACC-3′ primer sets having EcoR1 and SalI site respectively. All these constructs were verified by restriction digestion and subsequently by sequencing as described earlier^21^. Using the same set of primers for cloning the mutants in pSGFP2C1, TRPV1-WT, TRPV1-Arg575Asp, TRPV1-Arg575Asp/Asp576Arg were cloned into pmCherryC1 vector (Takara) and were verified by restriction digestion and sequencing.

## Primers used for Site Directed Mutagenesis

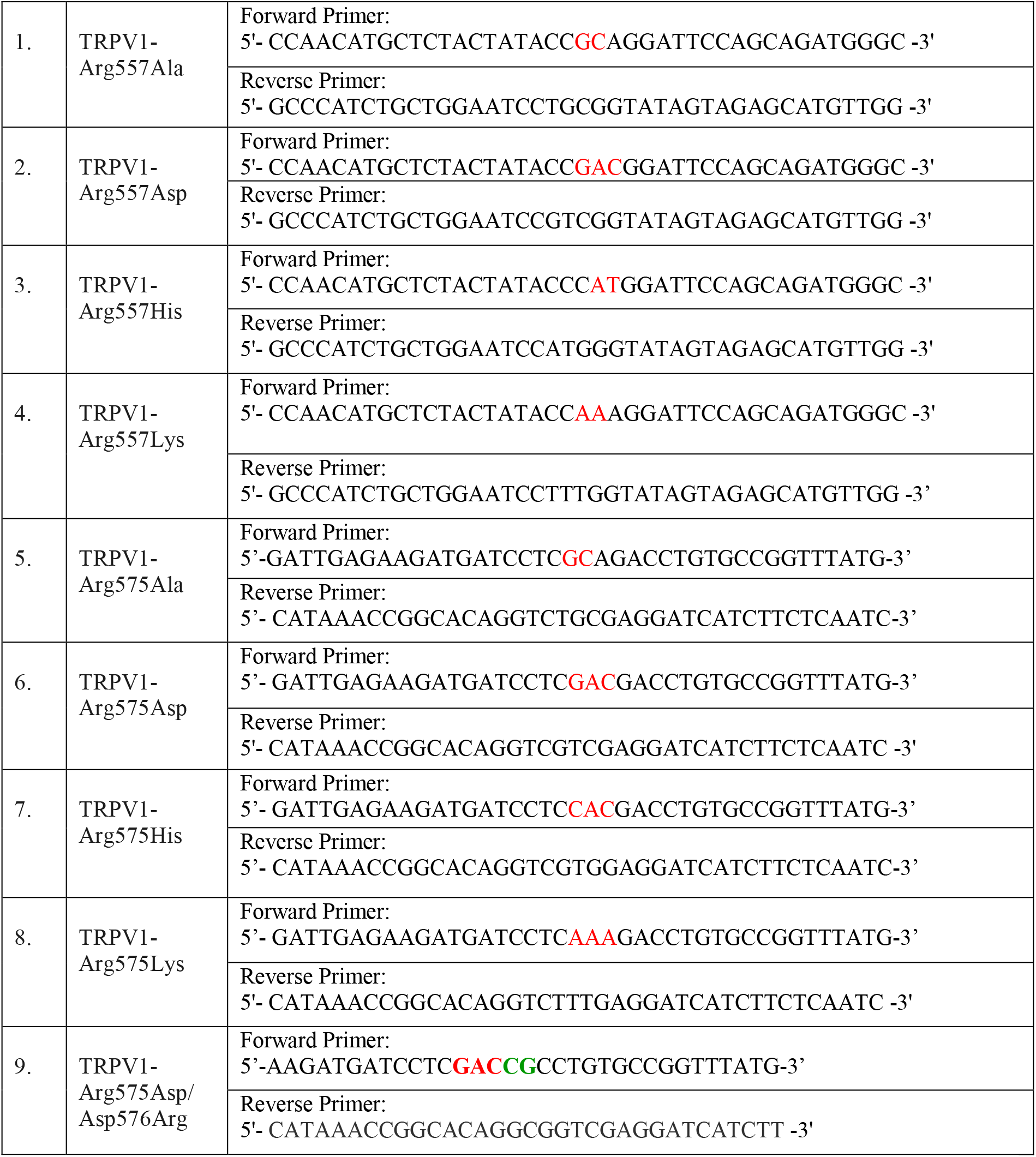

### Cell culture, transfection and imaging

F-11 cells were cultured in nutrient mixture F-12 Ham media supplemented with 10% Fetal Bovine Serum, L-glutamine (2mM), Streptomycin and Penicillin (100 mg/ml each, HiMedia, Bangalore, India) as described before^21^. Cells were maintained in a humidified atmosphere at 5% CO2 and 37°C. F-11 cells were transiently transfected with TRPV1-WT-GFP, all LWI mutants and Flotillin1-RFP (for co-localization studies) using Lipofectamine (Invitrogen). Co-localization experiments were repeated with Caveolin 1-RFP also. Cells were fixed with 4% PFA 36 hours post transfection. Fixed cells were then washed twice with 1X PBS (Himedia), treated with DAPI (Invitrogen, 1:1000 dilution) and then mounted using Flouromount-G (Southern Biotech) as described earlier^21^. In some experiments, F-11 cells expressing TRPV1-WT-GFP or LWI mutants were treated with 5mM Methyl β-Cyclodextrin (MβCD, Sigma) for 30 minutes for cholesterol depletion and subsequently fixed with 4% PFA. Untreated or MβCD-treated cells were stained with Cholera Toxin-B-594 (CTXB-594, Invitrogen, 1:200 dilution). The B subunit of *Vibrio cholera* i.e. Cholera Toxin is known to bind to the GM1 ganglioside present in plasma membrane and thus labels lipid raft^57^. Cells were subsequently observed using Zeiss LSM-800 or Olympus FV3000 Confocal Microscope.

Length, breadth, area and perimeter of F-11 cells transfected with TRPV1-WT-GFP, TRPV1-Arg575Asp-GFP and TRPV1-Arg575Asp/Asp576Arg was measured using Fiji Software. An ROI was drawn demarcating the periphery of each transfected cell and then using the tools of this software the various parameters were evaluated. The same was performed for F-11 cells transfected with TRPV1-WT-GFP and TRPV1-Arg575Asp-GFP with and without 5’-IRTX (1μM, Tocris).

### Cell labelling with Rhod-3 AM dye

F-11 cells were transiently transfected with TRPV1-WT-, TRPV1-Arg575Asp and TRPV1-Arg575Asp/Asp576Arg constructs (all in pSGFP2C1 vector) or only with pSGFP2C1 empty plasmid using Lipofectamine (Invitrogen). 24 hours post-transfection TRPV1-Arg575Asp-GFP and TRPV1-Arg575Asp/Asp576Arg-GFP dishes were incubated with Rhod-3 AM dye for different time points (10 min to 2 hours) and at different concentrations (up to 5 μM). TRPV1-WT-pSGFP2C1 dish was incubated with Rhod-3 AM dye with different concentrations (up to 5 μM) for different time points (15-60 minutes) and only pSGFP2C1 dish was incubated with 10 μM Rhod-3 AM dye for 60 minutes. After incubation the coverslips were washed with pre-warmed F-12 Ham’s complete media to remove unbound dye. Live cell imaging of only transfected cells was performed using Olympus FV3000 Confocal Microscope. Cells transfected with various constructs were tested for the amount of Rhod-3 AM dye uptake and thus imaged at different laser powers (0.1 to 1%) using solid state laser (ex/em = ~560/600 nm).

### Ca^2+^-imaging

F-11 cells were cultured in nutrient mixture F-12 Ham media supplemented with 10% Fetal Bovine Serum, L-glutamine (2 mM), Streptomycin and Penicillin (100 mg/ml each, HiMedia, Bangalore, India). Cells were maintained in a humidified atmosphere at 5% CO2 and 37°C as mentioned before. F-11 cells were transiently transfected with TRPV1-WT-pmCherryC1, TRPV1-Arg575Asp-pmCherryC1, TRPV1-Arg575Asp/Asp576Arg-pmCherryC1 and pmCherryC1. Each of these were co-transfected with the ultrasensitive protein calcium sensor pGP-CMV-GCaMP6f (Addgene)^58^. Approximately 24 hours after transfection, doubly transfected cells were imaged for 200 frames whereby cells were activated by adding 10 μM Capsaicin (Sigma) at the 30^th^ frame and again with 2 μM Ionomycin (Invitrogen) at the 120^th^ frame. Snaps of doubly transfected cells were acquired using 488 nm and 561 nm laser prior drug addition. Time series of live cells were conducted using 0.4% 488 nm laser and images were acquired for total 217 sec at a rate of 1 frame/1.085 secs.

### Quantification of Ca^2+^-imaging

Change in intensity of the ultrasensitive protein calcium sensor pGP-CMV-GCaMP6f was quantified over time throughout 200 frames. Each cell was considered to be a single ROI and then the change in intensity was quantified using Fiji software. The initial value was considered to be 1 and accordingly the changes were calculated and the graphs were plotted using GraphPad Prism 7 software. For analysing the Area Under Curve (AUC), the intensities of each construct were considered from 1-120 frames i.e. just before addition of Ionomycin. Using GraphPad Prism 7, AUC of each transfected cell were calculated and the statistical significance was assessed as described below. The time point of acquisition of highest intensity by each cell for a particular construct was analysed from the 30^th^ frame (point of addition of Capsaicin) to the frame where it acquired the highest intensity (this range was from 30-120 frame i.e. before addition of Ionomycin).

### Statistical test

Length, breadth, area and perimeter of transfected cells were calculated as described by Fiji software. Column graphs having just one variable were plotted for each construct from these values using GraphPad Prism 7. Statistical significance was analysed using One-way ANOVA (and non-parametric) which compared the mean of each column with the mean of every other column and the P-values were assessed using Tukey’s multiple comparison test. **_**The same procedure was followed for estimating the statistical significance of the calcium imaging plots for AUC and time point of highest intensity. The Pearson co-efficient (r) values were also calculated using GraphPad Prism 7 software.

## Acknowledgements

Intramural funding from NISER, Bhubaneswar is appreciated. Support and intellectual input from all the lab members is appreciated.

## Contributions

CG conceived the idea and designed all the experiments. RD1, DV, NT, RD2 performed all *in silico* analysis. RD1, DV, NT, AK and CG analysed all the *in silico* results. SS performed all the construct preparation, cell culture, sample preparation, all the microscopic experiments, image processing, quantification and their statistical analysis. AT performed all the Rhod-3-AM dye uptake experiments. CG wrote the paper. The model has been prepared by CG and SS All authors contributed towards manuscript editing.

## Corresponding author

<u>Chandan Goswami</u>.

## Ethics declarations

This manuscript has been prepared following highest level of scientific ethics concerned.

## Supplementary figures

**Figure S1:**
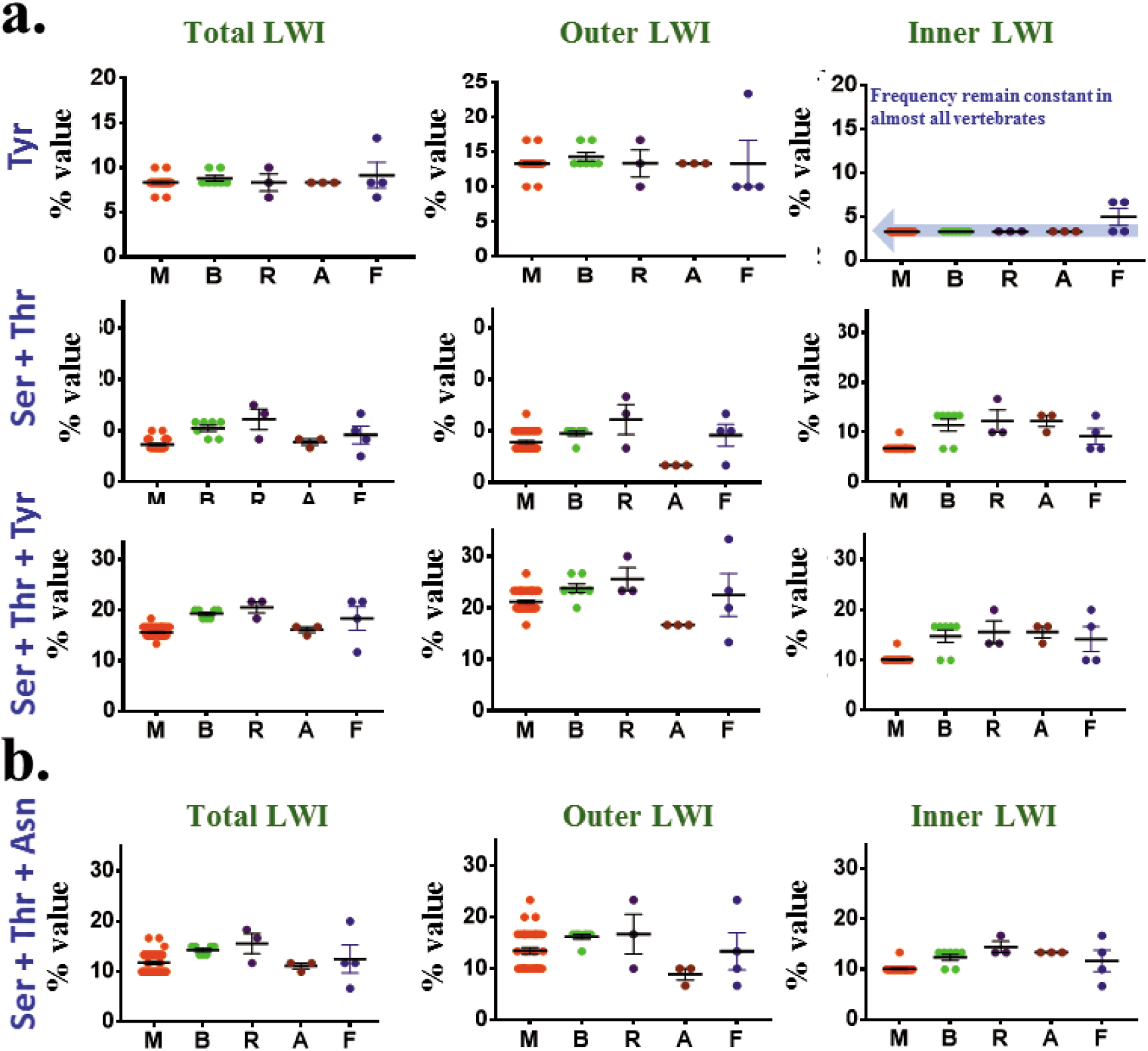
Some but not all post-translational modifications of TRPV1 may impose selection pressure on the amino acids located at lipid-water-interface. **a.** Shown are the frequency-of-occurrence of Tyr, Ser + Thr or all these three amino acids in lipid-water-interface. The frequency-of-occurrence of Tyr in the inner LWI is conserved in all vertebrates (indicated by a blue arrow as background) suggesting that Tyrosine-Kinase mediated phosphorylation events may play crucial role in the molecular selection of TRPV1 structure-function. Frequency-of-occurrence of Ser residues in the inner LWI region remain variable, yet positively selected in early vertebrates. Thr residue is negatively selected in all vertebrates and totally excluded in higher vertebrates suggesting that presence of Thr residues at the inner LWI does not provide adaptive advantages to all vertebrates. Therefore, Ser-Thr kinase-mediated regulations of TRPV as reported in several reports are likely to be species-specific and/or tissue specific events. **b.** Frequency-of-occurrence of Asp (which undergoes N-linked glycosylation), Ser and Thr (which undergo O-linked glycosylation) are shown. As the values representing Frequency-of-occurrence of individual as well as these three amino acids remain variable, especially in the outer LWI region, it is unlikely that specific glycosylation plays important role in the molecular selection of TRPV1. Different glycosylation’s reported in TRPV1 may therefore suggest events that are mostly tissue and species specific.

**Figure S2.**
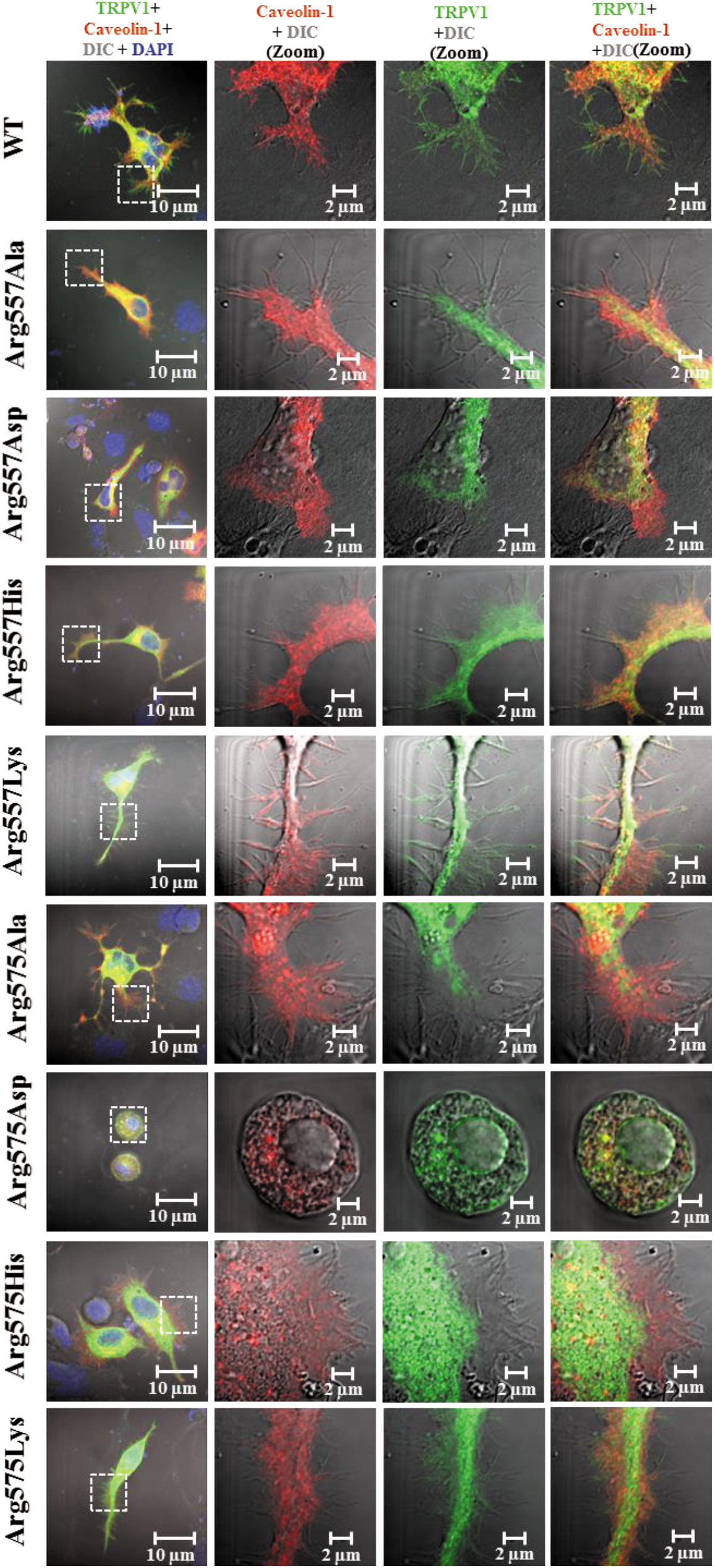
TRPV1-WT but not the Lipid Water Interface (LWI) mutants co-localize with overexpressed lipid raft markers. GFP-tagged (green) TRPV1 wild type (WT) and different LWI mutants were co-expressed with lipid raft marker Caveolin1-RFP (red) in F11 cells. Cells were fixed 36 hours post transfection and images were acquired by confocal microscope. TRPV1-WT shows distinct co-localization with Caveolin1-RFP in the membranous region. Notably, the LWI mutants are distinctly excluded from Caveolin1-RFP enriched membranous regions even after over expressing both.

**Figure S3.**
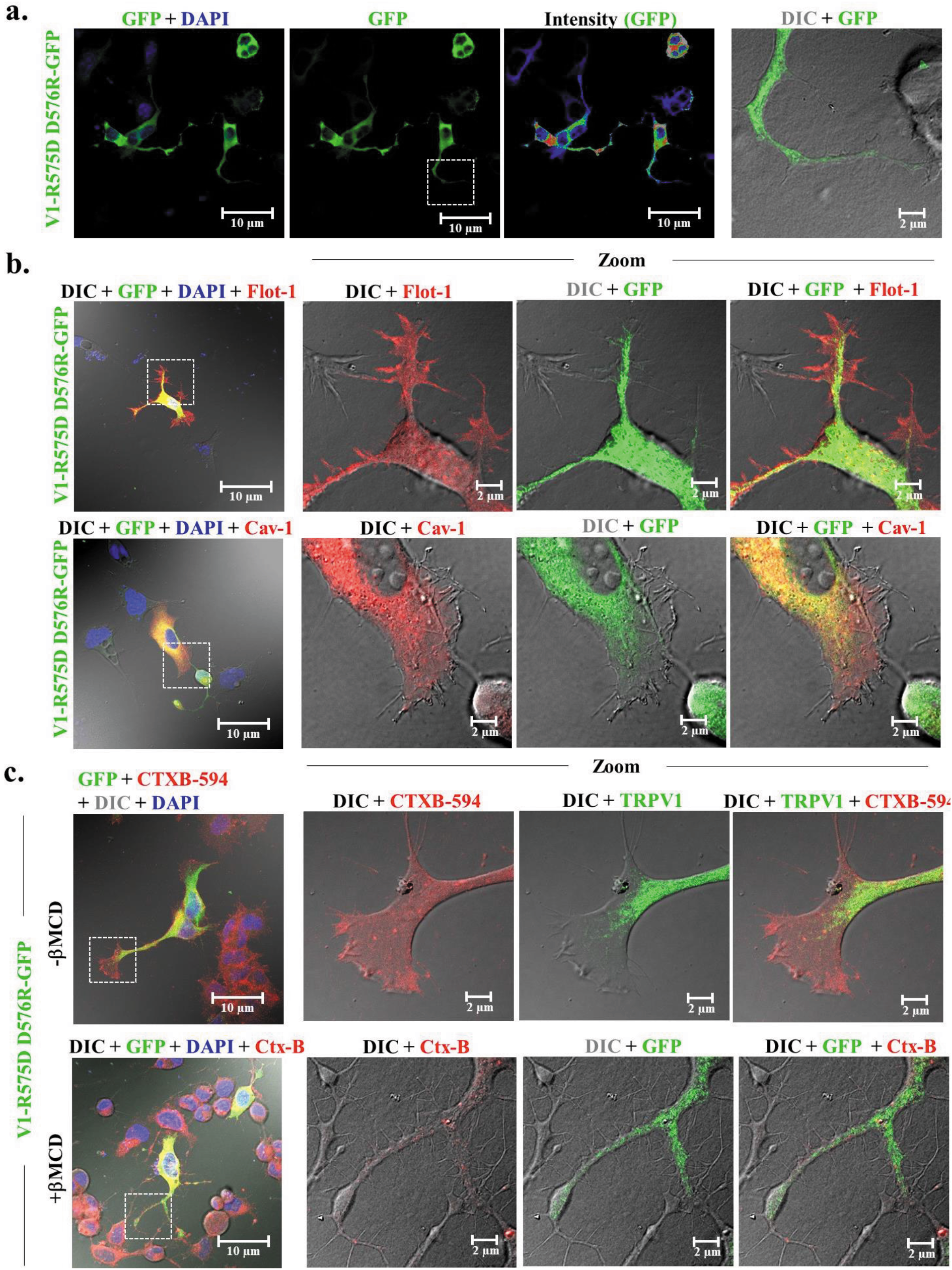
Localization of TRPV1-Arg575Asp/Asp576Arg-GFP in F-11 cells and its co-localization with different lipid raft markers. TRPV1-Arg575Asp/Asp576Arg in GFP vector was transiently transfected into F-11 cells. **a.** TRPV1-Arg575Asp/Asp576Arg fails to show distinct membrane localization. **b.** Overexpression of TRPV1-Arg575Asp/Asp576Arg-GFP with lipid raft markers Flotillin 1-RFP (Flot-1) and Caveolin 1-RFP (Cav-1). **c.** Localization of TRPV1-Arg575Asp/Asp576Arg-GFP with endogenous lipid raft marker CTXB-594 in control as well as cholesterol reduced condition (with 5 mM MβCD).

**Figure S4.**
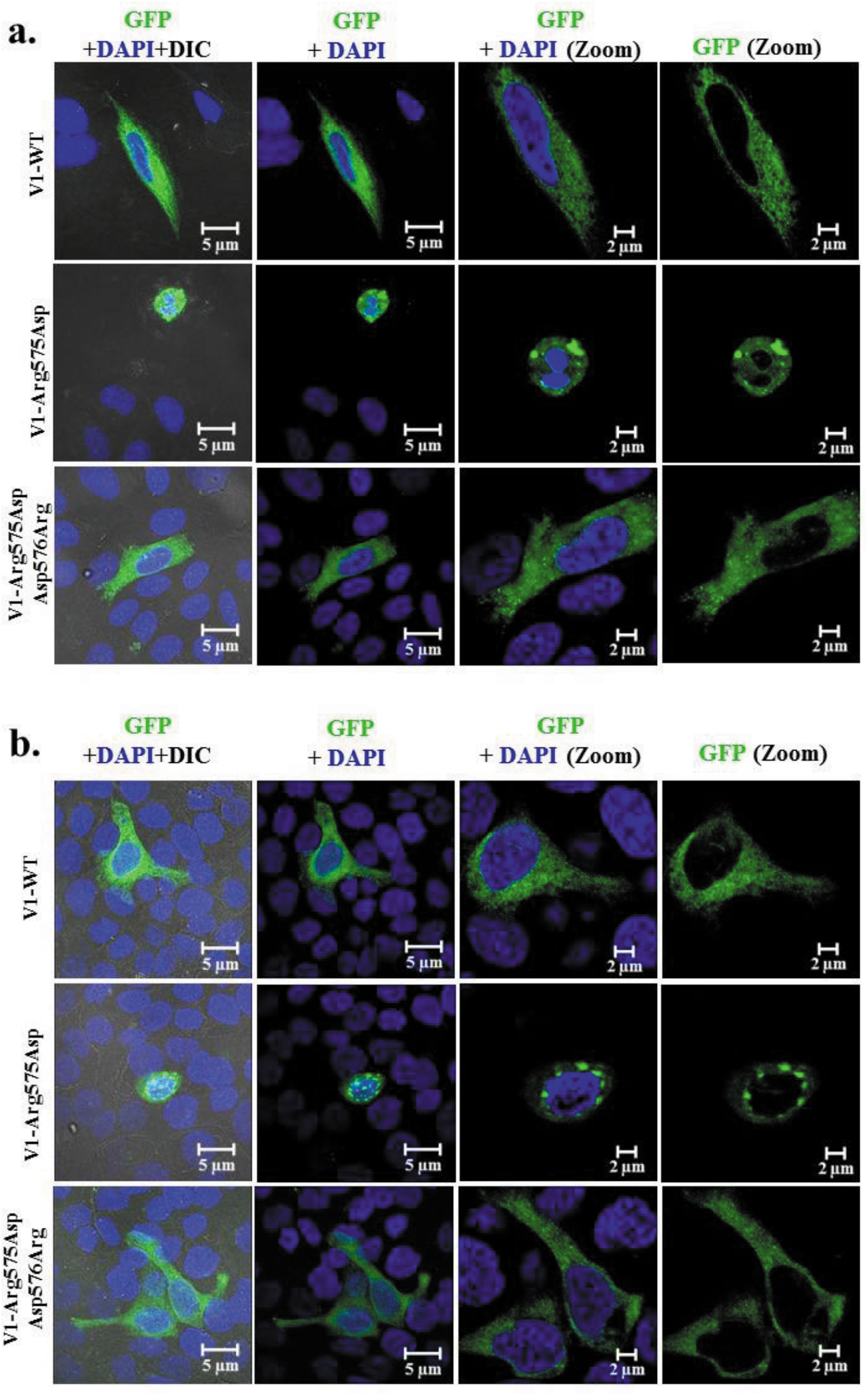
TRPV1-Arg575Asp induced lethality in SaOS and HaCaT cell lines can be restored by TRPV1-Arg575Asp/Asp576Arg. **a.** SaOS cells transiently transfected with TRPV1-WT, TRPV1-Arg575Asp and TRPV1-Arg575Asp/Asp576Arg (all in pSGFP2C1 vector). **b.** HaCaT cells transiently transfected with TRPV1-WT, TRPV1-Arg575Asp and TRPV1-Arg575Asp/Asp576Arg (all in pSGFP2C1 vector). In both cases lethality induced by TRPV1-Arg575Asp is restored by TRPV1-Arg575Asp/Asp576Arg.

**Figure S5.**
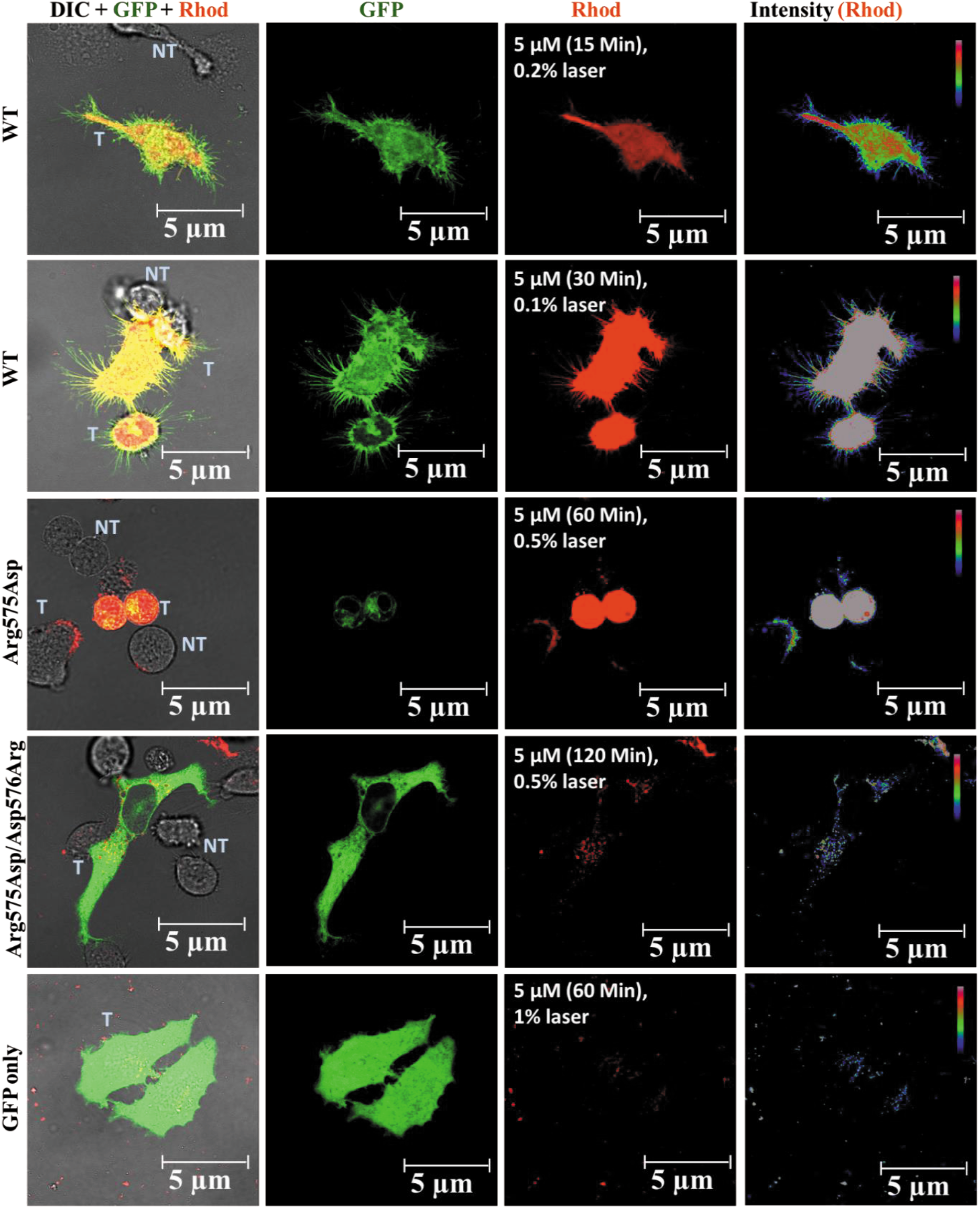
F11 cells expressing TRPV1-WT or TRPV1-Arg-575-Asp or TRPV1-Arg-575-Asp/Asp-576-Arg show differential uptake of Rhod-3AM dye. Cells were loaded with different amount of Rhod-3 AM dye for differ time points and imaged at different laser power. TRPV1-Arg-575-Asp/Asp-576-Arg uptakes much lesser dye than TRPV1-Arg-575-Asp mutant. The intensity of Rhod-3 AM dye is shown in pseudo colour (right most side).

**Figure S6.**
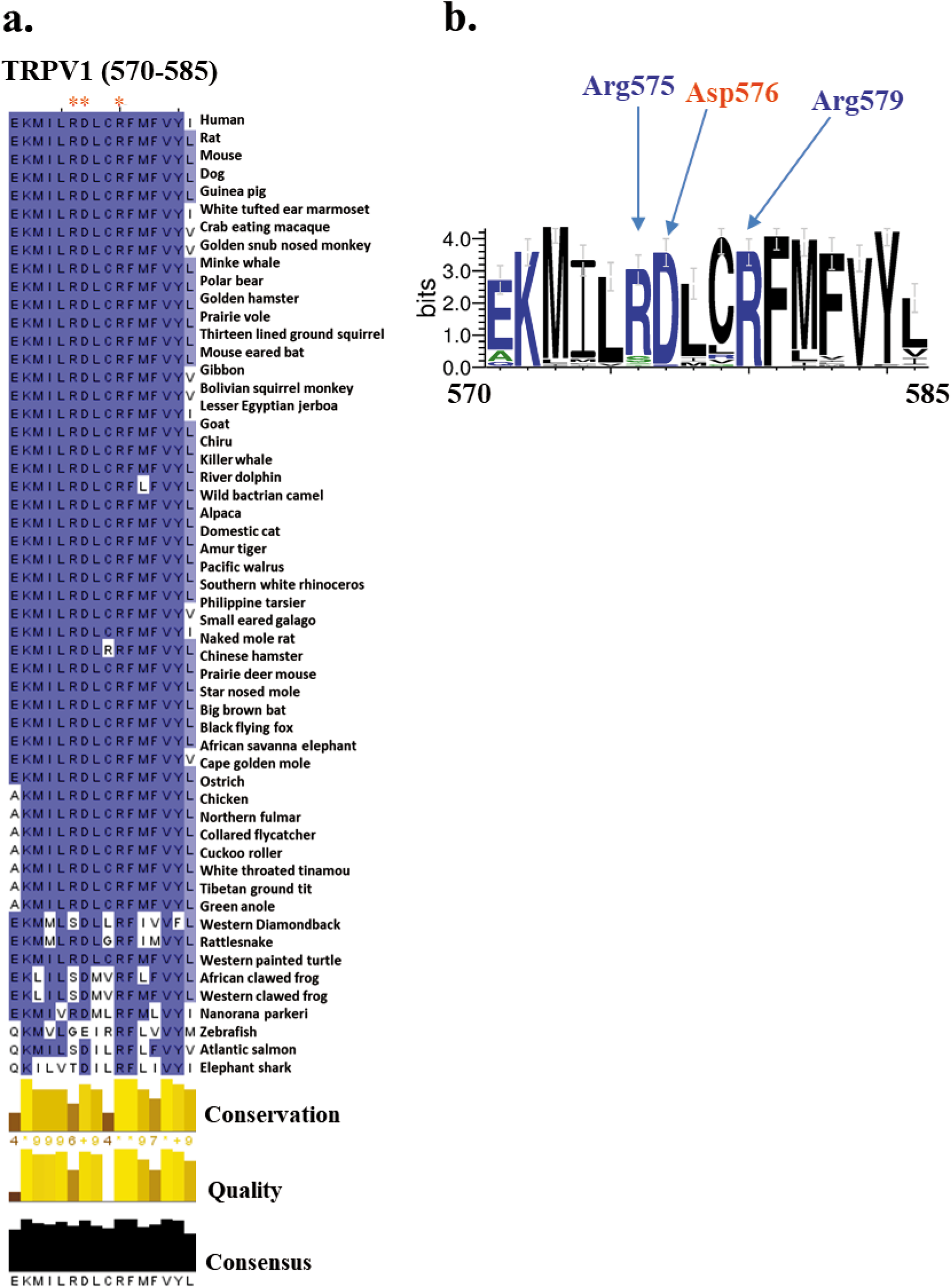
Conservation analysis of Arg575, Asp576 and Arg579 in TRPV1 across vertebrate evolution. **a.** Sequence alignment of TRPV1 amino acid (AA) stretch 570-585 across different vertebrate species are shown. Arg575, Asp576 and Arg579 have been highlighted (red star mark). **b.** SeqLogo of TRPV1 amino acid (AA) stretch 570-585 across different vertebrate species is shown. Arg575, Asp576 and Arg579 are indicated by arrows.

## References

1. Jentsch, T. J., Hübner, C. A. & Fuhrmann, J. C. Ion channels: Function unravelled by dysfunction. Nature Cell Biology (2004). doi:10.1038/ncb1104-1039

2. Yi, B. A., Minor, D. L., Lin, Y. F., Jan, Y. N. & Jan, L. Y. Controlling potassium channel activities: Interplay between the membrane and intracellular factors. Proc. Natl. Acad. Sci. U. S. A. (2001). doi:10.1073/pnas.191351798

3. Moreau, A., Gosselin-Badaroudine, P. & Chahine, M. Biophysics, pathophysiology, and pharmacology of ion channel gating pores. Frontiers in Pharmacology (2014). doi:10.3389/fphar.2014.00053

4. Chowdhury, S., Jarecki, B. W. & Chanda, B. A molecular framework for temperature-dependent gating of ion channels. Cell (2014). doi:10.1016/j.cell.2014.07.026

5. Hempling, H. G. Intracellular water and the regulation of cell volume and pH. Princ. Med. Biol. 4, 217–246 (1995).

6. Ciardo, M. G. & Ferrer-Montiel, A. Lipids as central modulators of sensory TRP channels. Biochimica et Biophysica Acta - Biomembranes (2017). doi:10.1016/j.bbamem.2017.04.012

7. Zhou, H. X. & McCammon, J. A. The gates of ion channels and enzymes. Trends in Biochemical Sciences (2010). doi:10.1016/j.tibs.2009.10.007

8. Kim, J.-B. Channelopathies. Korean J. Pediatr. 57, 1 (2014).

9. Imbrici, P. et al. Therapeutic approaches to genetic ion channelopathies and perspectives in drug discovery. Frontiers in Pharmacology (2016). doi:10.3389/fphar.2016.00121

10. Smutzer, G. & Devassy, R. K. Integrating TRPV1 Receptor Function with Capsaicin Psychophysics. Advances in Pharmacological Sciences (2016). doi:10.1155/2016/1512457

11. Elokely, K. et al. Understanding TRPV1 activation by ligands: Insights from the binding modes of capsaicin and resiniferatoxin. Proc. Natl. Acad. Sci. U. S. A. (2016). doi:10.1073/pnas.1517288113

12. Okamoto, N. et al. Effect of single-nucleotide polymorphisms in TRPV1 on burning pain and capsaicin sensitivity in Japanese adults. Mol. Pain (2018). doi:10.1177/1744806918804439

13. Caterina, M. J. et al. Impaired nociception and pain sensation in mice lacking the capsaicin receptor. Science (80-.). (2000). doi:10.1126/science.288.5464.306

14. Ghosh, A., Kaur, N., Kumar, A. & Goswami, C. Why individual thermo sensation and pain perception varies? Clue of disruptive mutations in TRPVs from 2504 human genome data. Channels (2016). doi:10.1080/19336950.2016.1162365

15. Sardar, P., Kumar, A., Bhandari, A. & Goswami, C. Conservation of tubulin-binding sequences in TRPV1 throughout evolution. PLoS One (2012). doi:10.1371/journal.pone.0031448

16. De Toni, L. et al. Heat sensing receptor TRPV1 is a mediator of thermotaxis in human spermatozoa. PLoS One (2016). doi:10.1371/journal.pone.0167622

17. Storozhuk, M. V., Moroz, O. F. & Zholos, A. V. Multifunctional TRPV1 Ion Channels in Physiology and Pathology with Focus on the Brain, Vasculature, and Some Visceral Systems. BioMed Research International (2019). doi:10.1155/2019/5806321

18. Liao, M., Cao, E., Julius, D. & Cheng, Y. Structure of the TRPV1 ion channel determined by electron cryo-microscopy. Nature (2013). doi:10.1038/nature12822

19. Chugunov, A. O., Volynsky, P. E., Krylov, N. A., Nolde, D. E. & Efremov, R. G. Temperature-sensitive gating of TRPV1 channel as probed by atomistic simulations of its trans- and juxtamembrane domains. Sci. Rep. (2016). doi:10.1038/srep33112

20. Morales-Lázaro, S. L. & Rosenbaum, T. Cholesterol as a Key Molecule That Regulates TRPV1 Channel Function. Adv. Exp. Med. Biol. 1135, 105–117 (2019).

21. Saha, S. et al. Preferential selection of Arginine at the lipid–water-interface of TRPV1 during vertebrate evolution correlates with its snorkeling behaviour and cholesterol interaction. Sci. Rep. (2017). doi:10.1038/s41598-017-16780-w

22. Wimley, W. C. & White, S. H. Experimentally determined hydrophobicity scale for proteins at membrane interfaces. Nature Structural Biology (1996). doi:10.1038/nsb1096-842

23. Goswami, C. et al. TRPV1 acts as a synaptic protein and regulates vesicle recycling. J. Cell Sci. (2010). doi:10.1242/jcs.065144

24. Hellwig, N. et al. TRPV1 acts as proton channel to induce acidification in nociceptive neurons. J. Biol. Chem. (2004). doi:10.1074/jbc.M402966200

25. Meyers, J. R. et al. Lighting up the senses: FM1-43 loading of sensory cells through nonselective ion channels. J. Neurosci. (2003). doi:10.1523/jneurosci.23-10-04054.2003

26. Binshtok, A. M., Bean, B. P. & Woolf, C. J. Inhibition of nociceptors by TRPV1-mediated entry of impermeant sodium channel blockers. Nature (2007). doi:10.1038/nature06191

27. Jansson, E. T. et al. Effect of cholesterol depletion on the pore dilation of TRPV1. Mol. Pain (2013). doi:10.1186/1744-8069-9-1

28. Verma, P., Kumar, A. & Goswami, C. TRPV4-mediated channelopathies. Channels (2010). doi:10.4161/chan.4.4.12905

29. Morales-Lázaro, S. L., Lemus, L. & Rosenbaum, T. Regulation of thermoTRPs by lipids. Temperature (2017). doi:10.1080/23328940.2016.1254136

30. Prescott, E. D. & Julius, D. A modular PIP2 binding site as a determinant of capsaicin receptor sensitivity. Science (80-.). (2003). doi:10.1126/science.1083646

31. Liu, M., Huang, W., Wu, D. & Priestley, J. V. TRPV1, but not P2X, requires cholesterol for its function and membrane expression in rat nociceptors. Eur. J. Neurosci. (2006). doi:10.1111/j.1460-9568.2006.04889.x

32. Sághy, É. et al. Evidence for the role of lipid rafts and sphingomyelin in Ca2+-gating of Transient Receptor Potential channels in trigeminal sensory neurons and peripheral nerve terminals. Pharmacol. Res. 100, 101–116 (2015).

33. Picazo-Juárez, G. et al. Identification of a binding motif in the S5 helix that confers cholesterol sensitivity to the TRPV1 ion channel. J. Biol. Chem. (2011). doi:10.1074/jbc.M111.237537

34. Das, R. & Goswami, C. TRPV4 expresses in bone cell lineages and TRPV4-R616Q mutant causing Brachyolmia in human reveals “loss-of-interaction” with cholesterol. Biochem. Biophys. Res. Commun. (2019). doi:10.1016/j.bbrc.2019.07.042

35. Munns, C. H., Chung, M. K., Sanchez, Y. E., Amzel, L. M. & Caterina, M. J. Role of the outer pore domain in transient receptor potential vanilloid 1 dynamic permeability to large cations. J. Biol. Chem. (2015). doi:10.1074/jbc.M114.597435

36. Kumari, S. et al. Influence of membrane cholesterol in the molecular evolution and functional regulation of TRPV4. Biochem. Biophys. Res. Commun. 456, 312–319 (2015).

37. MacCallum, J. L., Drew Bennett, W. F. & Peter Tieleman, D. Distribution of amino acids in a lipid bilayer from computer simulations. Biophys. J. (2008). doi:10.1529/biophysj.107.112805

38. Lee, A.. Lipid-protein interactions in biological membranes: a structural perspective. Biochim. Biophys. Acta - Biomembr. (2003). doi:10.1016/S0005-2736(03)00056-7

39. Strandberg, E. & Killian, J. A. Snorkeling of lysine side chains in transmembrane helices: How easy can it get? FEBS Lett. (2003). doi:10.1016/S0014-5793(03)00475-7

40. Payandeh, J., Pfoh, R. & Pai, E. F. The structure and regulation of magnesium selective ion channels. Biochimica et Biophysica Acta - Biomembranes (2013). doi:10.1016/j.bbamem.2013.08.002

41. Chan, D. I., Prenner, E. J. & Vogel, H. J. Tryptophan- and arginine-rich antimicrobial peptides: Structures and mechanisms of action. Biochimica et Biophysica Acta - Biomembranes (2006). doi:10.1016/j.bbamem.2006.04.006

42. Herce, H. D. et al. Arginine-rich peptides destabilize the plasma membrane, consistent with a pore formation translocation mechanism of cell-penetrating peptides. Biophys. J. (2009). doi:10.1016/j.bpj.2009.05.066

43. Allolio, C. et al. Arginine-rich cell-penetrating peptides induce membrane multilamellarity and subsequently enter via formation of a fusion pore. Proc. Natl. Acad. Sci. U. S. A. (2018). doi:10.1073/pnas.1811520115

44. Hristova, K. & Wimley, W. C. A look at arginine in membranes. Journal of Membrane Biology (2011). doi:10.1007/s00232-010-9323-9

45. Fantini, J. & Barrantes, F. J. How cholesterol interacts with membrane proteins: An exploration of cholesterol-binding sites including CRAC, CARC, and tilted domains. Frontiers in Physiology (2013). doi:10.3389/fphys.2013.00031

46. Caputo, G. A. & London, E. Cumulative effects of amino acid substitutions and hydrophobic mismatch upon the transmembrane stability and conformation of hydrophobic α-helices. Biochemistry (2003). doi:10.1021/bi026697d

47. Killian, J. A. & Von Heijne, G. How proteins adapt to a membrane-water interface. Trends in Biochemical Sciences (2000). doi:10.1016/S0968-0004(00)01626-1

48. Iyer, B. R., Vetal, P. V., Noordeen, H., Zadafiya, P. & Mahalakshmi, R. Salvaging the Thermodynamic Destabilization of Interface Histidine in Transmembrane β-Barrels. Biochemistry (2018). doi:10.1021/acs.biochem.8b00805

49. Sparks, K. A. et al. Comparisons of interfacial phe, tyr, and trp residues as determinants of orientation and dynamics for GWALP transmembrane peptides. Biochemistry (2014). doi:10.1021/bi500439x

50. Zheng, W. & Wen, H. Heat activation mechanism of TRPV1: New insights from molecular dynamics simulation. Temperature (2019). doi:10.1080/23328940.2019.1578634

51. Wen, H. & Zheng, W. Decrypting the Heat Activation Mechanism of TRPV1 Channel by Molecular Dynamics Simulation. Biophys. J. (2018). doi:10.1016/j.bpj.2017.10.034

52. Díaz-Franulic, I., Caceres-Molina, J., Sepulveda, R. V., Gonzalez-Nilo, F. & Latorre, R. Structure-driven pharmacology of transient receptor potential channel vanilloid 1. Molecular Pharmacology (2016). doi:10.1124/mol.116.104430

53. Numazaki, M., Tominaga, T., Toyooka, H. & Tominaga, M. Direct phosphorylation of capsaicin receptor VR1 by protein kinase Cε and identification of two target serine residues. J. Biol. Chem. (2002). doi:10.1074/jbc.C200104200

54. Zhang, F. et al. Heat activation is intrinsic to the pore domain of TRPV1. Proc. Natl. Acad. Sci. U. S. A. (2017). doi:10.1073/pnas.1717192115

55. Vyklický, L. et al. Calcium-dependent desensitization of vanilloid receptor TRPV1: A mechanism possibly involved in analgesia induced by topical application of capsaicin. Physiological Research (2008).

56. Valentine, M. L., Cardenas, A. E., Elber, R. & Baiz, C. R. Physiological Calcium Concentrations Slow Dynamics at the Lipid-Water Interface. Biophys. J. (2018). doi:10.1016/j.bpj.2018.08.044

57. Day, C. A. & Kenworthy, A. K. Functions of cholera toxin B-subunit as a raft cross-linker. Essays Biochem. (2015). doi:10.1042/bse0570135

58. Chen, T. W. et al. Ultrasensitive fluorescent proteins for imaging neuronal activity. Nature 499, 295–300 (2013).

